# Toward neuroanatomical and cognitive foundations of macaque social tolerance grades

**DOI:** 10.1101/2025.03.12.642838

**Authors:** S. Silvère, J. Lamy, C. Po, M. Legrand, J. Sallet, S. Ballesta

**Affiliations:** Laboratoire de Neurosciences Cognitives et Adaptatives, UMR 7364, Strasbourg, France; Centre de Primatologie de l’Université de Strasbourg, Niederhausbergen, France; ICube (UMR 7357), Université de Strasbourg-CNRS, Strasbourg, France; Univ Lyon, Université Lyon, Inserm, Stem Cell and Brain Research Institute, U1208, Bron, France; Wellcome Center for Neuroimaging, Dpt of Experimental Psychology, University of Oxford, Oxford, UK

**Keywords:** non-human primate, social tolerance, social brain, amygdala, MRI, comparative neuroanatomy

## Abstract

The macaque genus includes 25 species with diverse social systems, ranging from low to high social tolerance grades. Such interspecific behavioral variability provides a unique model to tackle the evolutionary foundation of primate social brain. Yet, the neuroanatomical correlates of these social tolerance grades remain unknown. To address this question, we expressed social tolerance grades within a novel cognitive framework and analyzed *post-mortem* structural scans from 12 macaque species. Our results show that amygdala volume is a subcortical predictor of macaques’ social tolerance, with high tolerance species exhibiting larger amygdala than low tolerance ones. We further investigated the developmental trajectory of amygdala across social grades and found that intolerant species showed a gradual increase in relative amygdala volume across the lifespan. Unexpectedly, tolerant species exhibited a decrease in relative amygdala volume across the lifespan, contrasting with the age-related increase observed in intolerant species—a developmental pattern previously undescribed in primates. Taken together, these findings provide valuable insights into the cognitive, neuroanatomical and evolutionary basis of primates’ social behaviors.

## Introduction

Complex social environment implies a greater cognitive demand of social representations and interactions, which is one of the driving forces behind the evolution of the primate brain (Dunbar, 2009; Freeberg et al., 2012; Heldstab et al., 2022). Correlations between social environment and variations in brain structures volumes have been reported, both in humans (Kanai et al., 2012; Maguire et al., 2000; Parkinson et al., 2017) and in non-human primates (NHP) (Noonan et al., 2014a; Sallet et al., 2011; Meguerditchian et al., 2021; Testard et al., 2022). In rhesus macaques (*M. mulatta*), previous studies have demonstrated that interindividual variation in social characteristics—such as hierarchical status (Noonan et al., 2014b) or group size (Sallet et al., 2011; Testard et al., 2022) – is associated with grey matter volume in core regions of the social brain, including the amygdala, the hippocampus, the superior temporal sulcus (STS), and the rostral prefrontal cortex (rPFC). Supporting the broader relevance of these findings across *Cercopithecinae*, a study in olive baboons (*Papio anubis*) revealed that individuals living in larger social groups exhibited greater total brain volumes, with an effect primarily driven by white matter (Meguerditchian et al., 2021).

Despite the existence of 25 species within the *Macaca* genus (Cooper et al., 2022; Cords, 2012; Ghosh et al., 2022; Thierry, 2007; Thierry et al., 2004), most neuroscience research focuses on two species, *M. mulatta* and *M. fascicularis* (and in rare cases *M. nemestrina* and *M. fuscata* (Carlo et al., 2010; Isa et al., 2009; Maranesi et al., 2014)). In spite of the relatively short evolutionary divergence time within this genus (6 to 8 million years (Perelman et al., 2011)), the various macaque species display a considerable interspecific variety of social behaviors while usually maintaining a multi-male, multi-female, and multi-generational social structure (Balasubramaniam et al., 2017; Thierry, 2007, 2000). These behavioral differences are characterized by different styles of dominance (Balasubramaniam et al., 2012), severity of agonistic interactions (Duboscq et al., 2014), nepotism (Berman and Thierry, 2010; Duboscq et al., 2013; Sueur et al., 2011) and submission signals (De Waal and Luttrell, 1985; Rincon et al., 2023), among the 18 covariant behavioral traits described in Thierry’s classification of social tolerance (Thierry, 2021, 2017, 2000).

Despite this large behavioral variability, macaque species display broadly similar general cognitive abilities (ManyPrimates et al., 2022). Specific differences observed in domains such as inhibitory control or social flexibility are thus more likely to reflect adaptive responses to species-specific social constraints, rather than intrinsic disparities in overall intelligence (Joly et al., 2017; Loyant et al., 2023). Altogether, the socio-behavioral diversity within the *Macaca* genus provides a compelling model to investigate how social ecology shapes cognition and its neural substrates.

The concept of social tolerance, central to this comparative approach, has sometimes been used in a vague or unidimensional way. As Bernard Thierry (2021) pointed out, the notion was initially constructed around variations in agonistic relationships – dominance, aggressiveness, appeasement or reconciliation behaviors – before being expanded to include affiliative behaviors, allomaternal care or male–male interactions (Thierry, 2021). These traits do not necessarily align along a single hierarchical axis but rather reflect a multidimensional complexity of social style, in which each trait may have co-evolved with others (Thierry, 2021, 2000; Thierry et al., 2004). Moreover, the lack of a standardized scientific definition has sometimes led to labeling species as “tolerant” or “intolerant” without explicit criteria (Gumert and Ho, 2008; Patzelt et al., 2014). These behavioral differences are characterized by different styles of dominance (Balasubramaniam et al., 2012), severity of agonistic interactions (Duboscq et al., 2014), nepotism (Berman and Thierry, 2010; Duboscq et al., 2013; Sueur et al., 2011) and submission signals (De Waal and Luttrell, 1985; Rincon et al., 2023), among the 18 covariant behavioral traits described in Thierry’s classification of social tolerance (Thierry, 2021, 2017, 2000).

To ground the investigation of social tolerance in a comparative neuroanatomical framework, we introduced a tentative working model that articulates behavioral traits, cognitive dimensions, and their potential subcortical neural substrates. Drawing upon 18 behavioral traits identified in Thierry’s comparative analyses (Thierry, 2021, 2007), we organized these traits into three core dimensions: socio-cognitive demands, behavioral inhibition, and the predictability of the social environment (Table 1). This conceptualization did not aim to redefine social tolerance itself, but rather to provide a structured basis for testing neuroanatomical hypotheses related to the volume of relevant subcortical areas and social style variability. It echoes recent efforts to bridge behavioral ecology and cognitive neuroscience by linking specific mental abilities – such as executive functions or metacognition – with distinct prefrontal regions shaped by social and ecological pressures (Bouret et al., 2024; Testard 2022).

**Table 1:** Cognitive and neuroanatomical categorization of behavioral traits associated with macaque social tolerance.

| Social trait | Underlying social consequences | Cognitive dimension | Neural correlate |
| --- | --- | --- | --- |
| Rate of reconciliation | Demands in recognizing partners, recalling social history and regulate affiliation<sup>1</sup>. | Higher socio-cognitive demands | Amygdala volume higher<sup>7,8,9</sup>;<br>Hippocampus volume higher<sup>9,10,11</sup> |
| Level of complexity of the communication system | Demands in interpreting social signals and adjusting communication to context<sup>2</sup>. |  |  |
| Development of male-to-male coalitions | Demands in social knowledge and strategic social decision<sup>3,4</sup>. |  |  |
| Development of cooperative behaviors | Demands in understanding intentions and coordinating actions during interactions<sup>5,6</sup>. |  |  |
| Intensity of aggression | Demands in inhibiting impulsive behaviors and regulating emotional responses<sup>12,13</sup>. | Better inhibitory control | Amygdala volume lower<sup>9,17</sup>;<br>Hippocampus volume unchanged<sup>17</sup>. |
| Degree of confidence of social play | Demands in adjusting behavior and inhibiting responses in mutual interactions<sup>14,15</sup>. |  |  |
| Resource distribution evenness | Demands in regulating emotions and adjusting behavior during competitive interactions<sup>16</sup>. |  |  |
| Kin bias (nepotism) | Kin knowledge is less informative to predict social relationships<sup>18,19</sup>. | Lower predictability of social environment | Amygdala volume higher<sup>7,8,9,17</sup>;<br>Hippocampus volume lower<sup>29,30,31</sup>. |
| Degree of dominance asymmetry | Conflicts are not always won by dominant individuals, making social outcomes less predictable<sup>16,20</sup>. |  |  |
| Development of formal submission signals | Social communication during conflict is less informative to predict conflict outcomes<sup>21,22</sup>. |  |  |
| Intensity of female rank inheritance | Matrilinear knowledge is less informative to predict social relationship<sup>23,24</sup>. |  |  |
| Rate of affiliative contact | Affiliative network is denser and therefore harder to predict<sup>25,26,27</sup>. |  |  |
| Rate of counter-aggression | Subordinates are more likely to retaliate making social outcomes less predictable<sup>3,28</sup>. |  |  |
| Rate of immature interference in mating | Mounting behaviour trigger more social interactions which induce 'erratic' interaction<sup>14</sup>. |  |  |
| Degree of mother protectiveness | Recognizing individuals and anticipating social risks in interaction<sup>32</sup>. | Unclassified | / |
| Development of allomothering behavior | Understanding social roles and interpreting socio-affective cues<sup>33</sup>. |  |  |
| Delayed male dispersal | Navigating hierarchies through flexible behavior and inhibition<sup>16</sup>. |  |  |
| Centrality of top-ranking males | Tracking relationships and managing distributed social networks<sup>34</sup>. |  |  |

1. Thierry and Sapolsky, 2000; 2. Liebal et al., 2014; 3. Petit et al., 1997; 4. Silk, 1999; 5. Demaria and Thierry, 2001; 6. Micheletta et al., 2012; 7. Bickart et al., 2011; 8. Sallet et al., 2011; 9. Testard et al., 2022; 10. Kanai et al., 2012; 11. Todorov et al., 2019; 12. Adams et al., 2015; 13. Loyant et al., 2023; 14. Petit et al., 2008; 15. Scopa and Palagi, 2016; 16. Thierry, 2007; 17. Tottenham et al., 2010; 18. Silk, 2002; 19. Silk et al., 2003; 20. de Vries et al., 2006; 21. Flack et al., 2006; 22. Waller et al., 2013; 23. Hill and Okayasu, 1995; 24. Kutsukake, 2000; 25. Duboscq et al., 2013; 26. Massen et al., 2010; 27. Silk et al., 2003; 28. Balasubramaniam et al., 2012; 29. Kim et al., 2015; 30. Lyons et al., 2001; 31. Meyer and Hamel, 2014; 32. Maestripieri, 1994; 33. Fairbanks, 1990; 34. Sueur et al., 2011;

Navigating social life in primate societies requires substantial cognitive resources: individuals must not only track multiple relationships, but also regulate their own behavior, anticipate others’ reactions, and adapt flexibly to changing social contexts. Taken advantage of databases of magnetic resonance imaging (MRI) structural scans, we conducted the first comparative study integrating neuroanatomical data and social behavioral data from closely related primate species of the same genus to address the following questions: To what extent can differences in volumes of subcortical brain structures be correlated with varying degrees of social tolerance? Additionally, we explored whether these dispositions reflect primarily innate features, shaped by evolutionary processes, or acquired through socialization within more or less tolerant social environments.

The first category, socio-cognitive demands, refers to the cognitive resources needed to process, monitor, and flexibly adapt to complex social environments. Linking those parameters to neurological data is at the core of the social brain theory (Dunbar, 2009). Macaques’ social systems require advanced abilities in social memory, perspective-taking, and partner evaluation (Freeberg et al., 2012). This is particularly true in tolerant species, where the increased frequency and diversity of interactions may amplify the demands on cognitive tracking and flexibility. Tolerant macaque species typically live in larger groups with high interaction frequencies, low nepotism, and a wider range of affiliative and cooperative behaviors, including reconciliation, coalition-building, and signal flexibility (Thierry, 2021, 2000). Tolerant macaque species also exhibit a more diverse and flexible vocal and facial repertoire than intolerants ones which may help reduce ambiguity and facilitate coordination in dense social networks (Rincon et al., 2023; Scopa and Palagi, 2016; Rebout 2020). Experimental studies further show that macaques can use facial expressions to anticipate the likely outcomes of social interactions, suggesting a predictive function of facial signals in managing uncertainty (Micheletta et al., 2012; Waller et al., 2016). Even within less tolerant species, like *M. mulatta*, individual variation in facial expressivity has been linked to increased centrality in social networks and greater group cohesion, pointing to the adaptive value of expressive signaling across social styles (Whitehouse et al., 2024).

The second category, inhibitory control, includes traits that involve regulating impulsivity, aggression, or inappropriate responses during social interactions. Tolerant macaques have been shown to perform better in tasks requiring behavioral inhibition and also express lower aggression and emotional reactivity than intolerants macaques both in experimental and in natural contexts (Joly et al., 2017; Loyant et al., 2023). These features point to stronger self-regulation capacities in species with egalitarian or less rigid hierarchies. More broadly, inhibition – especially in its strategic form (self-control) – has been proposed to play a key role in the cohesion of stable social groups. Comparative analyses across mammals suggest that this capacity has evolved primarily in anthropoid primates, where social bonds require individuals to suppress immediate impulses in favour of longer-term group stability (Dunbar and Shultz, 2025). This view echoes the conjecture of Passingham and Wise (2012), who proposed that the expansion of lateral prefrontal area BA10 in anthropoids enabled the kind of behavioural flexibility needed to navigate complex social environments (Passingham et al., 2012).

The third category, social environment predictability, reflects how structured and foreseeable social interactions are within a given society. In tolerant species, social interactions are more fluid and less kin-biased, leading to greater contextual variation and role flexibility, which likely imply a sustained level of social awareness. In fact, as suggested by recent research, such social uncertainty and prolonged incentives are reflected by stress-related physiology : tolerant macaques such as *M. tonkeana* display higher basal cortisol levels, which may be indicative of a chronic mobilization of attentional and regulatory resources to navigate less predictable social environments (Sadoughi et al., 2021).

Each behavioral trait was individually evaluated based on existing empirical literature regarding the types of cognitive operations it likely involves. When a primary cognitive dimension could be identified, the trait was assigned accordingly. However, some behaviors – such as maternal protection, allomaternal care, or delayed male dispersal – do not map neatly onto a single cognitive process. These traits likely emerge from complex configurations of affective and social-motivational systems, and may be better understood through frameworks such as attachment theory (Suomi, 2008), which emphasizes the integration of social bonding, emotional regulation, and contextual plasticity. While these dimensions fall beyond the scope of the present framework, they offer promising directions for future research, particularly in relation to the hypothalamic and limbic substrates of social and reproductive behavior.

Rather than forcing these traits into potentially misleading categories, we chose to leave them unclassified within our current cognitive framework. This decision reflects both a commitment to conceptual clarity and the recognition that some behaviors emerge from a convergence of cognitive demands that cannot be neatly isolated. This tripartite framework, leaving aside reproductive-related traits, provides a structured lens through which to link behavioral diversity to specific cognitive processes and generate neuroanatomical predictions.

We therefore associated these three categories with neuroanatomical hypotheses regarding variations in the volume of two subcortical structures of interest, the amygdala and the hippocampus. Based on existing literature on the effects of the socio-cognitive demands, inhibitory control and social unpredictability – particularly when it induces sustained activation of stress-related systems – on the volume of these two regions (Bickart et al., 2011; Caetano et al., 2021; Coplan et al., 2014; Haley et al., 2012; Howell et al., 2014; Lyons et al., 2001; Sallet et al., 2011), we hypothesized that increased cognitive demands from the social environment could lead to a differential effect on amygdala and hippocampus volumes (Bickart et al., 2011; Haley et al., 2012; Lyons et al., 2001; Sallet et al., 2011). We summarized our working hypotheses into a table comparing grade 4 and grade 1 species (Table 1).

This table summarizes the 18 behavioral traits used to characterize social tolerance grades in the *Macaca* genus, based on Thierry’s comparative framework. Each trait is associated with a description of its underlying cognitive implications and assigned—when applicable—to one of three cognitive dimensions: (i) socio-cognitive demands (e.g., tracking partners, coordinating actions), (ii) behavioral inhibition (e.g., regulating impulsivity), or (iii) predictability of the social environment (e.g., anticipating interaction outcomes). The final column presents the hypothesized effects of these dimensions on the volume of two subcortical structures: the amygdala and the hippocampus. Traits that could not be clearly assigned to a specific cognitive dimension—often related to maternal or reproductive strategies—are marked as “unclassified”. This framework is used to generate testable predictions about the neural substrates of social style diversity in macaques.

We tested our hypothesis using 43 *post-mortem* MRI acquisitions of 12 macaque species representing the four grades of social tolerance. The dataset was both composed of samples from open access databases (Milham et al., 2018; Navarrete et al., 2018; Sakai et al., 2018) as well as newly and unpublished samples from the collection of the Centre de Primatologie de l’Université de Strasbourg (CdP) and INSERM-Oxford University. These samples include brain images of *M. tonkeana* and *M. thibetana,* two macaque species that have never been scanned before as well as scan of *M. nigra* that is rare in the existing literature (Navarrete et al., 2018; Sakai et al., 2023). Up to this date, only one study has included tolerant species of macaque monkeys in such neuroanatomical comparative framework (Jones et al., 2021). While Jones et al. (2020) identified interspecific differences in amygdala microstructure and serotonergic innervation, their histological approach did not assess structural volumes at the whole-brain level. To our knowledge, our study is the first to report neuroanatomical correlates of social tolerance grades of the *Macaca* genus based on *post-mortem* MRI volumetric analysis. This approach reveals two key findings. First, across species, amygdala volume is positively correlated with social tolerance grades, with more tolerant macaque species exhibiting larger amygdala volumes. Second, developmental trajectories of the amygdala diverge according to social style: in intolerant species, amygdala volume increases with age – as commonly reported in the literature (Schumann et al., 2019) – whereas in tolerant species, we observe an unexpected marked decrease over the lifespan. This study offers a novel and valuable perspective by comparing inter-species brain structures to investigate the functioning of the social brain, while accounting for key socio-cognitive variables.

## Results

We obtained structural MRI scans of 43 macaques of 12 macaque species. Using a semi-automated registration to an atlas (SARM, Hartig et al., 2021), we extracted amygdala and hippocampus volumes and analyzed whether these covaried with social grade and age, using a Bayesian model. The raw relations between the main response variables (the amygdala’s and hippocampus volumes) are depicted in **Figure 1**.

**Figure 1:**
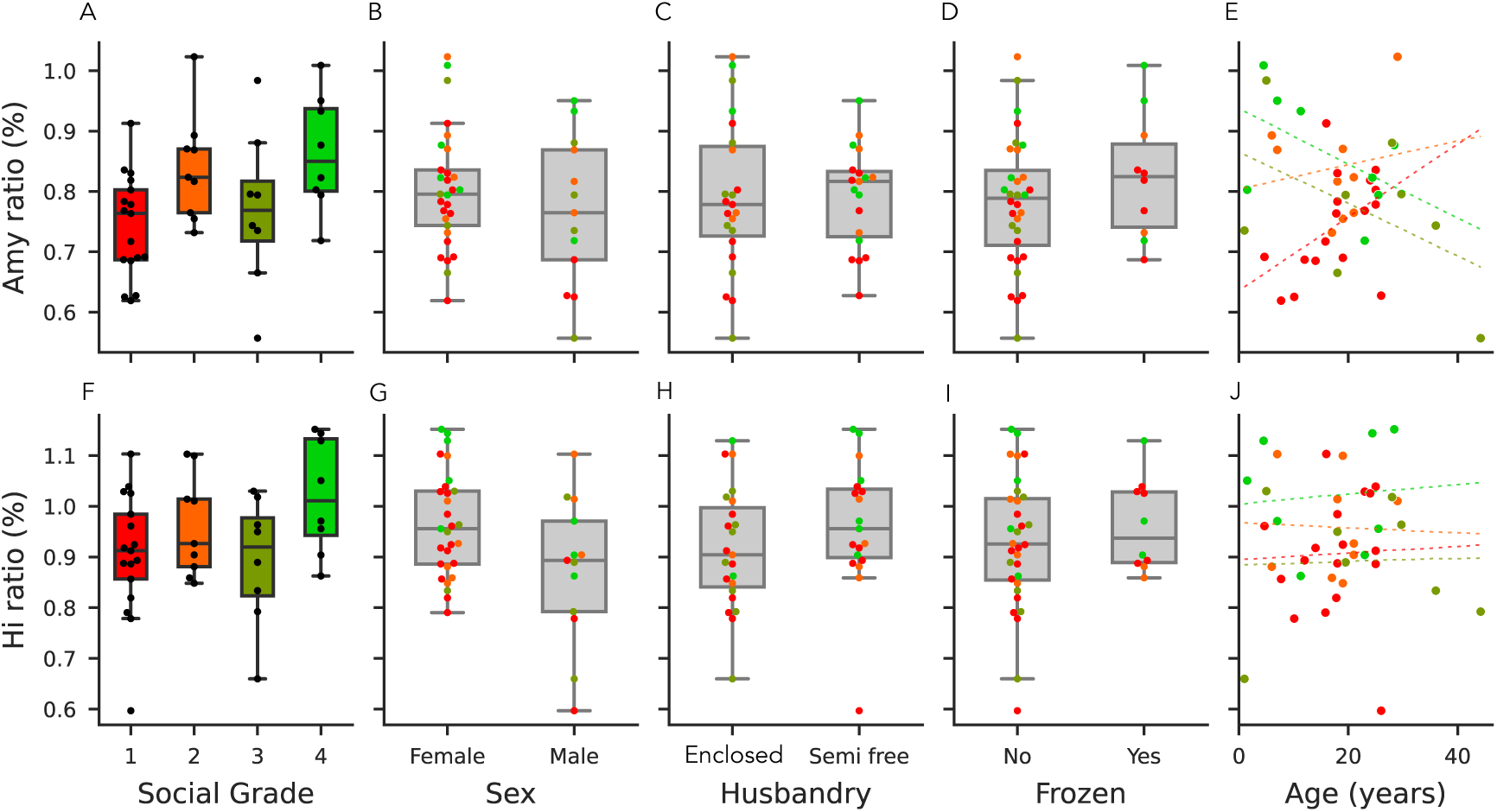
Model predictors of the amygdala and hippocampus, and volume predictions across social tolerance grades. First row **(A–D)**: Model predictors and responses for amygdala volume. The volume ratio is calculated as the amygdala volume divided by the total brain volume (excluding the myelencephalon and cerebellum). **(A)** Distribution of amygdala volume ratios across social tolerance grades. **(B)** Distribution of amygdala volume ratios by sex. **(C)** Distribution of amygdala volume ratios by husbandry condition (enclosed vs. semi-free). **(D)** Distribution of amygdala volume ratios by the frozen status. **(E)** Distribution of amygdala volume ratios by age. Second row **(F–J)**: Model predictors and responses for hippocampal volume. The volume ratio is calculated as the hippocampal volume divided by the total brain volume (excluding the myelencephalon and cerebellum). **(F)** Distribution of hippocampal volume ratios across social tolerance grades. **(G)** Distribution of hippocampal volume ratios by sex. **(H)** Distribution of hippocampal volume ratios by husbandry condition (enclosed vs. semi-free). **(I)** Distribution of hippocampal volume ratios by the frozen status. **(I)** Distribution of hippocampal volume ratios by age. Panels A-E et F-J share the same y-axis.

### Model quality and coefficients

The R² coefficient of determination of the model indicated a large proportion of variability accounted for by the model (90% credible interval: [0.87, 0.97]). The effect of sex was minimal for the amygdala but more pronounced for the hippocampus (**Figure 1, 2**), whereas husbandry had a limited effect on both regions of interest. Amygdala volume increased with social grade (independently of its interaction with age) and with age (independently of its interaction with social grade). However, the interaction between social grade and age suggested that the trajectory of amygdala volume over the lifespan differs across social grades, as detailed below. Total brain volume was included as a covariate in the model to account for interindividual differences in brain size. For descriptive purposes, its distribution across social grades is shown in Figure S1.”

**Figure 2:**
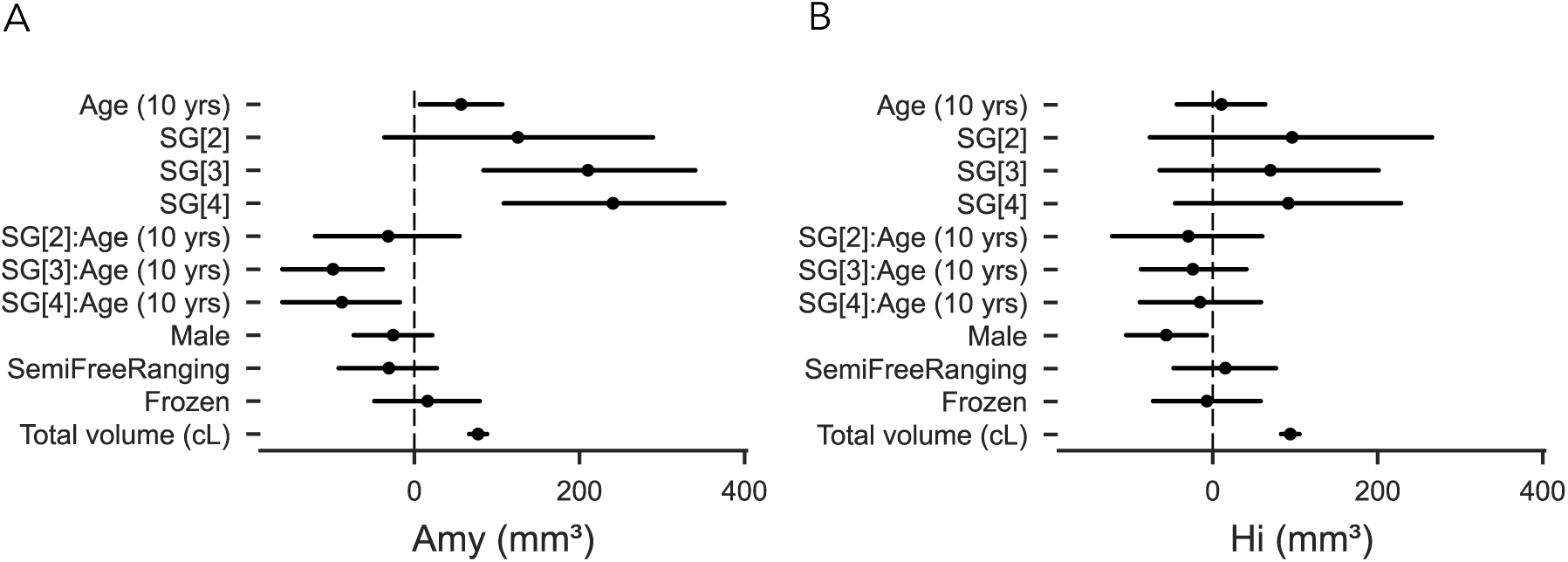
Parameters of the model. **(A)** Parameters of the model for the amygdala volume. **(B)** Parameters of the model for the hippocampal volume. SG [x]: Social Grade [x] vs Social Grade [1]; SG[x]:Age (10 years): Social Grade-Age interaction.

Despite limited sample size, the interaction between social grade and age suggested a differential trajectory of amygdala volume across the lifespan among different social grades (**Figure 1,2**).

To further assess group differences, we implemented Bayesian hypothesis testing using a Region of Practical Equivalence (ROPE, (Kruschke, 2015)) approach, with the ROPE defined as ±0.1×σ. This method allows classification of results into three categories: (a) a credible difference if the entire posterior interval lies outside the ROPE; (b) an absence of difference if it lies entirely within the ROPE; and (c) inconclusive if it overlaps the ROPE. For the amygdala, social grade 4 (SG4, i.e. tolerant) individuals had credibly larger volumes than social grade 1 (SG1, i.e. intolerant) individuals up to 19 years of age. For the hippocampus, the posterior distribution of the SG4–SG1 difference briefly exceeded the ROPE between approximately 13 and 18 years of age, indicating a credible difference in this age window. Outside this range, the intervals overlapped the ROPE, resulting in inconclusive evidence. However, the 90% posterior intervals remained entirely above zero at all ages, indicating that SG1 individuals never had larger hippocampal volumes than SG4 (**Figure 3**).

**Figure 3:**
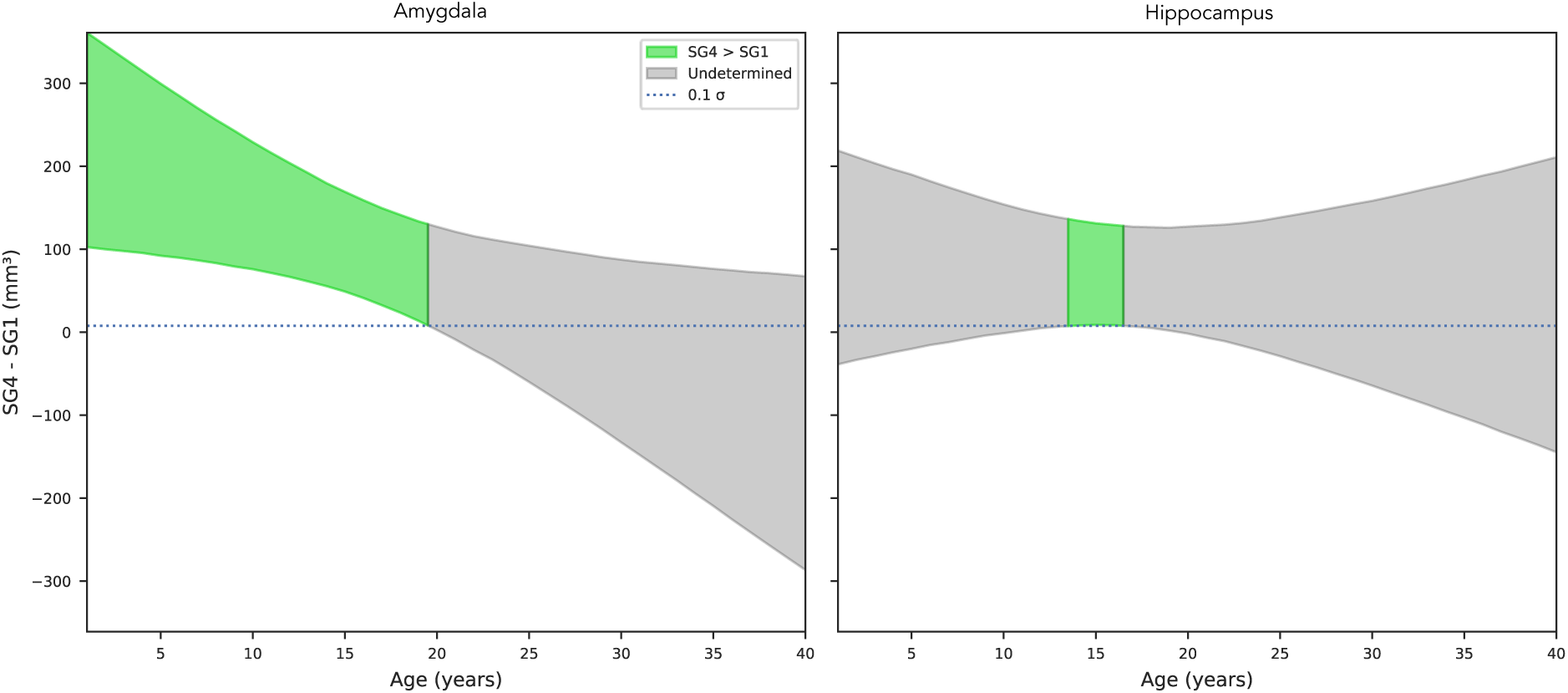
Bayesian hypothesis testing using a Region of Practical Equivalence (ROPE) to assess volume (in mm^3^) differences between Social Grade 4 (SG4; tolerant) and Social Grade 1 (SG1; intolerant) across age, for the amygdala (left) and hippocampus (right). Curves represent median posterior estimates, and shaded areas show 90% credible intervals. Gray bands indicate the ROPE (±0.1σ). For the amygdala, the difference is credible until ∼19 years. For the hippocampus, a credible effect is observed only between ∼13 and 18 years.

### Predicted data

Figure 4 illustrates how amygdala’s volume development varies with an individual’s social grade over their lifespan. Two results stand out: first, individuals in Social Grade 1 showed a distinct pattern of amygdala volume development compared to other social grades. Although Grade 1 individuals had a smaller amygdala volume in early years compared to the other Social Grades, the amygdala’s volume variation slope was steeper than for the other Grades (slope with 90% credible intervals [0.6, 11.0]). This increase contrasted with trends observed in Grades 3 (slope with 90% CI [−7.6, −0.9]) and 4 (slope with 90% CI [−8.0, 1.9]), which showed a decrease in volume over time. Individuals in Grade 2 also showed a slight increase in amygdala volume (slope with 90% CI [−4.4, 9.3]), similar to grade 1 but not as steep.

**Figure 4:**
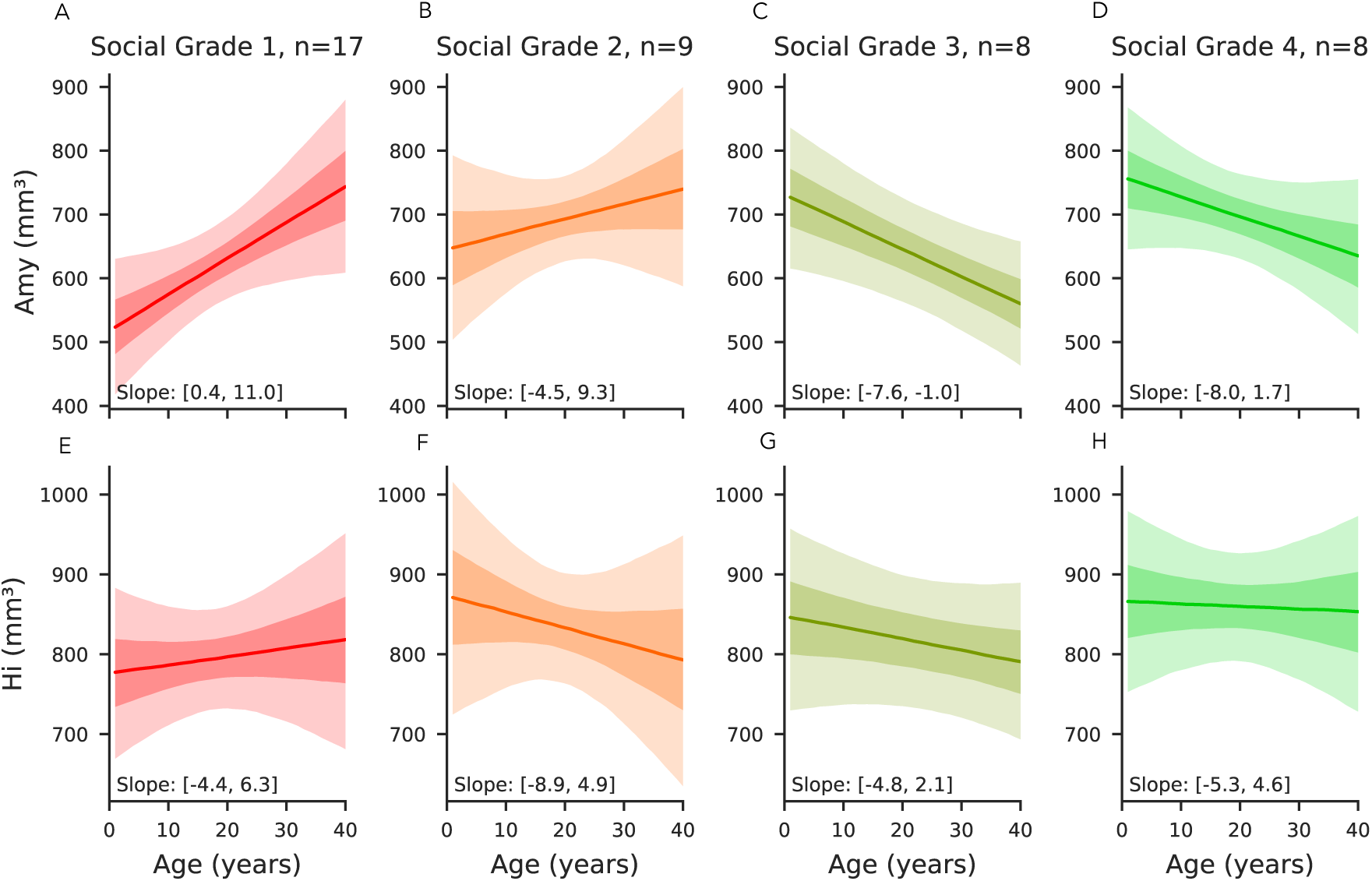
Volume predictions across social tolerance grades of the amygdala and hippocampus. All panels represent the predictions of the multivariate Bayesian linear model, where all the variables are kept constant (including total brain volumes) in order to represent the effect of age only on the volume of amygdala and hippocampus in mm^3^. First row **(A–D)**: Predicted amygdala volume across social tolerance grades over the lifespan. **(A)** Predicted amygdala volume as a function of age for grade 1 (intolerant) individuals. **(B)** Predicted amygdala volume as a function of age for grade 2 individuals. **(C)** Predicted amygdala volume as a function of age for grade 3 individuals. **(D)** Predicted amygdala volume as a function of age for grade 4 (tolerant) individuals. Second row **(E–H)**: Predicted hippocampal volume across social tolerance grades over the lifespan. **(E)** Predicted hippocampal volume as a function of age for grade 1 individuals. **(F)** Predicted hippocampal volume as a function of age for grade 2 individuals. **(G)** Predicted hippocampal volume as a function of age for grade 3 individuals. **(H)** Predicted hippocampal volume as a function of age for grade 4 individuals. In the plots, the solid lines represent the mean predicted values, and the shaded areas indicate the 90% credible intervals, with each social grade shown in a distinct color.

When comparing Grade 1 and Grade 4, individuals in Grade 4 showed larger amygdala volumes until approximately 19 years of age (Figure 4).

As expected from model predictions, hippocampal volume showed limited variation across age and social grades. ROPE-based hypothesis testing revealed that hippocampal volume in SG1 individuals was never greater than in SG4 individuals, supporting a consistent asymmetry in favor of more tolerant species.

## Discussion

We studied for the first time the neuroanatomical foundation of the naturally observed diversity of behavioral traits within the *Macaca* genus. We have assembled a unique database representing nearly half of the known macaque species, with a variety of ages, sexes, and origins. 12 species of them had never been scanned prior to our study. Our investigation focused on the subcortical structures of the brain and more especially the amygdala. This set of nuclei sometimes referred to as a hub of brain networks related to sociality and their social lives are well known for their roles in the stress response (Hölzel et al., 2010), emotional regulation and social cognition (Noonan et al., 2014b; Bickart et al., 2014; Amaral et al., 2003). Based on *post-mortem* MRI acquisitions from 12 of the 25 macaque species, we showed that amygdala volume correlated with the social tolerance grade and increased with the level of the social grade. Secondly, grade 4 species had a significantly higher amygdala volume at the start of their lives, which decreased over time, compared with grade 1 species, which showed the opposite trend. Finally, further hypothesis testing suggested that species of grade 1 never exhibited larger hippocampus and may have smaller hippocampus around age 15 when compared to those of grade 4. In accordance with our hypotheses (**Table 1**), our findings substantiated the assertions that (i) social tolerance is rooted in neuroanatomical differences that can be detected at an early stage of individuals’ development, (ii) social styles exert differential influence on subcortical structures throughout individuals’ lifespan and (iii) such phenomenon should be mainly driven by the socio-cognitive demands that vary with species social style (as evidenced by the higher amygdala and hippocampus volumes in higher tolerance species).

### A neuroanatomical account for social tolerance differences

The social tolerance grades are based on previous ethological observations of behaviors across different species of the genus (Thierry, 2021, 2007). From these observations, we identified three major cognitive processes—socio-cognitive demands, social predictability, and inhibitory control (see **Table 1**)—that underpin the observed behaviors. Among the behaviors we classified with “high social-cognitive demand”, several have been previously described in the literature as particularly discriminating between grades 1 and 4. It included greater social network density in grade 4 species (a consequence of, *inter alia*, low nepotism in tolerant species, facilitating interactions between unaffiliated conspecifics) (Balasubramaniam et al., 2017; Sueur et al., 2011), more complex facial mimics as well as a more complex communication system (Dobson, 2012; Rincon et al., 2023; Scopa and Palagi, 2016; Zannella et al., 2017), a significantly higher rate of reconciliation (Thierry, 2007), and a higher frequency of cooperative behaviors, including male-to-male coalition behaviors (De Waal and Luttrell, 1989; Thierry et al., 2008) in grade 4 species.

At first glance, ones may presume that species of lower social tolerance level, that displayed more overall aggressive behaviours, would have a larger amygdala when compared to more tolerant species. In fact, amygdala ablation or activity modulation in macaque monkeys showed that animals displayed less aggressive and different patterns of social behaviours (Amaral, 2002; Bliss-Moreau 2013, Wellman et al., 2016, Forcelli et al., 2016, Raper et al, 2019), supporting the idea that amygdala activity can promote aggressive behaviours. Our study revealed an opposite trends, amygdala was found to be larger in more tolerant species, and this apparent contradiction invites a more integrative view of the amygdala (Adolphs, 2009; Amaral et al., 2003; Pessoa, 2010), not only as a relay for emotional reactivity, but as a multifunctional hub embedded in complex social networks (Bickart et al., 2014). Such complex patterns of behavioural implications are also reflected at the cellular level, as amygdala is composed of several different nuclei that are broadly connected with other brain areas that may displayed opposite functions (Zikopoulos et al, 2017; Giacometti et al., 2023). Rather than opposing social cognition and emotion, our results support the view that emotional processing is deeply intertwined with social function—both being subserved by overlapping neural circuits (Domínguez-Borràs and Vuilleumier, 2022). It also suggests that additional neural mechanisms, particularly those involving prefrontal and anterior cingulate regions implicated in the top-down regulation of affect and social behavior, may contribute to shaping species differences in social tolerance (Ochsner and Gross, 2008). While our analysis compares social tolerance grades with variations in brain structure, the originality of our framework also lies in introducing three cognitive dimensions that bridge behavioral traits and neural substrates. This intermediate level of interpretation allows us to move beyond simple grade-to-structure associations, toward a more mechanistic understanding of the links between social behavior, cognition, and neuroanatomy.

Amygdala volume has also been shown to correlate positively with social network complexity in grade 1 species, as measured by the social network size of individuals (Sallet et al., 2011; Testard et al., 2022), or by the social status of the animals (Noonan et al., 2014b). This supports the idea that the amygdala is sensitive to both structural social features and dynamic aspects of social networks.

### Developmental trajectories and life-history plasticity

We are then led to question the origin of the social tolerance effect on amygdala volume, not in terms of a rigid nature versus nurture dichotomy, but in terms of differential developmental trajectories. Cross-fostering experiments (De Waal and Johanowicz, 1993), along with our own results, suggest that social tolerance grades reflect both early, possibly innate predispositions and later environmental shaping. Moreover, the behavioral shifts observed in cross-fostered individuals underscore the plasticity of social style acquisition and the role of early social environment in shaping neural substrates of social behavior. Notably, tolerant species exhibit larger amygdala volumes early in life, while intolerant species show a progressive increase across the lifespan—a pattern that suggests a dual influence of biological programming and cumulative social experience. These environmental influences likely arise from both species-specific social dynamics—such as variations in affiliative behavior and social play (Beltrán Francés et al., 2020)—and broader ecological conditions that structure the demands of social life. The age-related volumetric changes we observed, particularly the divergence in developmental trajectories between tolerant and intolerant species, reinforce this idea and echo previous reports of amygdala growth patterns in humans and macaques (Schumann et al., 2019; Uematsu et al., 2012). Taken together, these elements support the view that social tolerance is not fixed but emerges from the interplay between inherited developmental programs and the specific socio-ecological environments in which individuals mature.

Notably, the developmental trajectory of the amygdala in tolerant species does not align with that of intolerant species or with human developmental patterns (Schumann et al., 2019; Uematsu et al., 2012). This finding suggests that neurodevelopmental pathways may exhibit significant variation among phylogenetically closely related primate species, potentially serving as an effective evolutionary target for adapting socio-ecological behaviours to environmental demands. Moreover, we observed that in old individuals (typically above 19 years), relative amygdala volume in grade 1 species could match that of grade 4 species — despite being significantly smaller earlier in life. Due to a limited sample size of our study, this crossing trend, already accounted for by our continuous age model, should be further investigated. These results call for cautious interpretation of age-related variation and further emphasize the importance of longitudinal studies integrating both behavioral, cognitive and anatomical data in non-human primates, which would help to better understand the link between social environment and brain development (Song et al., 2021).

### Hippocampal Volume and Social Cognitive Demands in Tolerant Species

A credible difference in hippocampal volume favoring SG4 individuals was only revealed between approximately 13 and 18 years of age by our hypothesis testing using a ROPE framework. Outside this range, the difference remained overall positive but inconclusive. This restricted window of significance, along with the unidirectional trend across the lifespan, suggests that increased hippocampal volume may nonetheless be associated with higher social tolerance, at least adulthood. At first glance, this observation may appear to contrast with literature linking chronic stress to reduced hippocampal size (Kim et al., 2015; Lyons et al., 2007). However, as previously discussed, *M. tonkeana* (a high-tolerance species) combines elevated basal cortisol levels with a relatively large hippocampus (Sadoughi et al., 2021; Vandeleest et al., 2016), which suggests that glucocorticoid exposure alone does not account for hippocampal variation in this context. Instead, our findings are more consistent with the idea that hippocampal structure reflects species-specific cognitive demands associated with navigating complex and tolerant social environments—such as spatial memory, social recognition, or contextual learning (Han et al., 2021; Sallet et al., 2011). Within the conceptual framework introduced in this study, these results point to the importance of socio-cognitive requirements—rather than social environmental unpredictability or behavioral inhibition abilities—as potential drivers of interspecific variation in hippocampal anatomy. Comparative measurements and observations at individual level along with in vivo MRI from these same considered individuals may help to further understand how social tolerance can relate to cognitive abilities and its neural underpinning.

### Limits of the study and future directions

While our dataset is comprehensive in terms of the number of macaque species included certain limitations must be acknowledged. For instance, phylogenetic analyses were beyond the reach of this study and integrating these statistical approaches could clarify the extent to which interspecific differences in brain structure and social behavior are due to shared ancestry or convergent evolution (Ghosh et al., 2022; Heuer et al., 2025).

Although we explained some interspecies variability, adding subjects to our database will increase statistical power and will help addressing potential confounding factors such as age or sex in future studies. One will benefit from additional information about each subject. While considered in our modelling, the social living and husbandry conditions of the individuals in our dataset remain poorly documented. The living environment has been considered, and the size of social groups for certain individuals, particularly for individuals from the CdP, have been recorded. However, these social characteristics have not been determined for all individuals in the dataset. As previously stated, the social environment has a significant impact on the volumetry of certain regions. Furthermore, there is a lack of data regarding the hierarchy of the subjects under study and the stress they experience in accordance with their hierarchical rank and predictability of social outcomes position (McCowan et al., 2022). In addition, our treatment of sex differences was limited. Although sex was included as a covariate in the Bayesian models, the strong imbalance in our dataset—favoring females (2:1 ratio)—precludes robust conclusions about sex-specific trajectories. Some trends, particularly regarding hippocampal volume, suggest potential interactions between sex, age, and social grade, but these effects remain exploratory. Addressing them adequately would require larger and more balanced samples, along with behavioral or hormonal data to capture intra-sexual variability. It is therefore important to recognize that confirmation of our findings should be achieved by analyzing datasets in which all of these confounding factors can be controlled more effectively.

While our study identifies the amygdala as a key subcortical structure associated with interspecific variation in social tolerance, it is important to acknowledge several neuroanatomical limitations. First, our analyses were conducted on the amygdala as a whole, without distinction between its internal nuclei. Although we used the SARM atlas (Hartig et al., 2021), which offers a high-quality parcellation for *Macaca mulatta*, the precision of this template does not allow for fully reliable automatic segmentation of amygdala subnuclei across the diverse species included in our dataset. As a result, our volumetric measures may conflate distinct functional subregions, potentially masking more localized effects. In this context, histological approaches remain essential for characterizing fine-grained neuroanatomical differences, as illustrated by Jones et al. (2021), who reported interspecific variation in cell density and serotonergic innervation within the amygdala (Jones et al., 2021). Future studies combining MRI-based volumetry with *post-mortem* histology would allow more precise identification of which subregions underlie the observed differences in social tolerance.

### Cognitive and neural perspectives on our understanding of social tolerance

Future directions linking behavior, cognition and neuroanatomy could deepen our understanding of the roots of social tolerance among macaque species. This could lead to a better operationalization of the concept that could be applied to a wider range of non-human primate species. From a neural perspective, studying the cortical regions associated with social tolerance represents a promising yet ambitious goal. In fact, there is a variability within primate species in cerebral organization (Amiez et al., 2023, 2019) which is likely to be found across the *Macaca* genus. Considering this cerebral variability would require extensive efforts to properly assess interspecies differences, making it beyond the scope of the current study that focus on subcortical areas. However, as a starting point, exploring the connections between the amygdala, hippocampus, and medial prefrontal cortex could provide crucial insights into the neural correlates of social tolerance. These regions are central to stress regulation, socio-cognitive processing, and decision-making, all of which are likely impacted by social tolerance grades (Caetano et al., 2021; Coplan et al., 2014; Kim et al., 2015; Phelps and LeDoux, 2005; Sapolsky et al., 1990). In humans, repeated positive or stressful experiences have been demonstrated to alter the size of subcortical brain areas such as the hippocampus or amygdala (Davidson and McEwen, 2012) and impair neuroplasticity (Phelps, 2006). Neuronal plasticity and learning have been identified as contributing factors to variations in the ROI volume, including the amygdala and hippocampus, particularly in humans (Maguire et al., 2000; Taren et al., 2013). Additionally, our conceptual framework opens avenues for advanced neuroimaging techniques such as diffusion tensor imaging (DTI) (Howard et al., 2023; Zhang et al., 2013) or multiparametric MRI (Mulholland et al., 2024), which could be used to explore white matter connectivity or microstructural changes. Our findings also emphasize the need to develop individual-level measures of social tolerance (Dubuc et al., 2012; DeTroy et al., 2021). Fine-tuning these measures would allow more precise correlations between behavioral data and neuroanatomical features. By operationalizing the concept of social tolerance on cognitive dimensions, our work aims at enriching the framework through which primate sociality is currently studied.

## Conclusion

Our study provides novel insights into the relationship between amygdala volume and social tolerance in macaques, offering an innovative perspective on the neuroanatomical basis of social cognition. Using a comparative approach across 12 macaque species, we uncovered a revealing relationship: low-tolerance species start their life with a smaller amygdala compared to their socially tolerant counterparts. In addition, intolerant species show an increase in amygdala volume, whereas highly tolerant species show the opposite trend. These findings refine conventional views of the amygdala by highlighting its broader role in both emotional regulation and complex social cognition. The observed differences in amygdala volume with respect to social tolerance grades suggest that the development and plasticity of the amygdala seem to be intricately linked to the social environment and experiences of the species. Larger amygdala in socially tolerant species may reflect an enhanced capacity to process complex social information, facilitating better social interactions, cooperative behavior, and conflict management. Alternatively, the observed increase in amygdala volume in socially intolerant species over time may be explained by heightened socio-cognitive demands, rather than being solely attributed to chronic stress or emotional reactivity. While earlier studies emphasized the role of the amygdala in stress response, recent findings are in lines with our results which suggest that amygdala’s functions extend to broader aspects of social cognition. These findings have profound implications for our understanding of social brain evolution as well as underscoring the importance of developmental stage and the social environment being crucial drivers of neuroanatomical adaptations. In addition, although hippocampal volume showed less pronounced and more variable differences across social grades, a credible effect was observed between 13 and 18 years of age. Across all ages, SG4 individuals consistently exhibited larger hippocampal volumes than SG1, supporting the possibility that this region also contributes to social cognitive processes in tolerant species—especially during developmental phases associated with social maturation. This study, at the interface of primatology and cognitive neuroscience, also provides a framework for investigating the impact of the social environment on brain development and pave the way for future research to unravel the complexities of brain evolution and sociality.

## Material & Methods

### Brain specimen collection

To allow comprehensive cross-species comparisons in the *Macaca* genus, a dataset of 43 *post-mortem* specimens has been constituted through collaborations with multiple research centers, each contributing unique expertise and resources (**Table 2**). The collaborating institutions included **(Table S1)**:

The **Centre de Primatologie de l’Université de Strasbourg (CdP):** provided valuable brain data derived from 20 brain samples. Among those, one sample (*M. nigra*) was obtained as part of a collaboration with the zoo of Mulhouse (https://www.zoo-mulhouse.com/).

**Samples from INDI-PRIME-DE** (Milham et al., 2018)**: The Japan Monkey Center** provided 5 *post-mortem* MRI acquisitions to the dataset (Sakai et al., 2018). **Utrecht University:** contributed to the dataset with 13 *post-mortem* MRI acquisitions (Navarrete et al., 2018).

**INSERM-Oxford University:** 5 *post-mortem* MRI acquisitions came from this collaboration. This addition offered more variety of acquired data mostly in age and sex (Milham et al., 2018).

### Ethical considerations

The study was conducted in accordance with ethical guidelines and was approved by the ethical committee of the Centre de Primatologie de l’Université de Strasbourg which is authorized to house NHP (registration B6732636). The research further complied with the EU Directive 2010/63/EU for animal experiments. All subjects from the CdP died of natural or accidental cause; no macaque was euthanized in the sole frame of the project. These specimens originated from CdP, and their collection followed rigorous ethical considerations. The specimens were either obtained from previous collections—where full bodies were preserved in dedicated freezers—or from individuals of the CdP that had died from natural causes. The *post-mortem* MRI data from INSERM-Oxford University were acquired from deceased animals that died of causes unrelated to the present research project. As such, the research did not require a Home Office License as defined by the Animals (Scientific Procedures) Act 1986 of the United Kingdom.

### Brain extraction technique

*Post-mortem* MRI images acquisition of macaque brains is central to our study, more specifically, in translational studies of homologous brain regions. Brain extraction is a crucial process in neuroscience research for studying the internal brain structure of animals. Through the acquisition at the CdP of 20 *post-mortem* anatomical MRI scans of brains from six different species of macaques, we were able to refine a brain extraction technique - whether previously frozen or fresh - to minimize specimen handling artefacts and obtain image quality suitable for optimal use by the scientific community. The detailed extraction technique protocol established and used for our brain extractions is available in **Document S1**. Briefly, the head is reclined forward to expose the neck, muscles are removed to access the atlanto-occipital junction, which is then incised to allow head dislocation (**Figure S1**). An osteotome and hammer are used, ensuring no cerebellar herniation. The skull cap is carefully drilled using a rotary tool and removed (see **Figure S1. A, B** and **Table S2** for required tools), and the brain is extracted by severing the olfactory peduncles, internal carotid arteries, and cranial nerves. Specimens are then fixed in 10% buffered formaldehyde for 7 days (see **Figure S1. D.**) and in phosphate buffered saline (PBS) for 3-4 days before being placed in Fluorinert^TM^ for MRI acquisition, ensuring minimal air bubbles and optimal image quality (Sébille et al., 2019) see **Figure S2**).

### Sampling methods & measurements

Structural images were collected through both the open access databases and collaborations (Milham et al., 2018; Navarrete et al., 2018; Sakai et al., 2018), but also carried out at the IRIS platform of the ICube laboratory in Strasbourg for *post-mortem* samples kept at the CdP (see **Table S3** for the information relating to the acquisition of anatomical MRI images). The final dataset consists of 43 anatomical scans after pruning data with missing age or sex information (10 individuals), with both T_1_ and T_2_-weighted images. Due to their different origins, the images in the dataset did not follow the exact same acquisition protocols (different scanners and acquisition parameters, **Table 2**). In addition, *post-mortem* brain preservation and perfusion protocols are different, which may also influence the images obtained. Volume measurements were performed using a semi-automatic method to register individual images to the Subcortical Atlas of the Rhesus Macaque (SARM) (Hartig et al., 2021) **(Figure S1.E.)**. Due the large orientation discrepancies across the research centers, the images were first manually realigned (translation and rotation) with the atlas using ITK-SNAP (Yushkevich et al., 2006), then non-linearly registered using ANTs (Avants et al., 2011). The segmentation maps of the atlas were then transported to the subject space to extract the volume of the regions of interest. **Figure S1** details the image processing for volume extractions (see **Figure S2**). To ensure the accuracy of the SARM on our dataset, which includes 11 species other than *M. mulatta* (the species used for SARM development (Hartig et al., 2021)), we calculated the Dice Similarity Coefficient (DSC) (Zou et al., 2004). This was done by manually segmenting, using a tablet (Wacom Cintiq^®^ 16 and ITK SNAP software), the amygdala in each acquisition and comparing the overlapping voxels between the manual segmentation and the SARM segmentation. With a DSC of 0.96, we confirm the robust performance of the SARM across our entire dataset.

### Final dataset characteristics

The dataset is composed of 12 distinct macaque species with a total of 43 individual specimens for analysis. There is a strong sex imbalance with more females than males. The age range spans from 1 to 44.20 years, with an average age of 18.2 ± 9.4 years (standard deviation) (**Figure 5. B, D.)** with two outliers above 35 years old. Most importantly, based on our research question, the social grade distribution of our dataset (**Figure 5. A.)** is more represented by grade 1 than grade 4 species as these species are very rare in zoo or in research centers and most of them protected as endangered species. The MRI acquisitions from the 43 individuals were standardized to the NMT template (Jung et al., 2021). Data included amygdala or hippocampus volume and a computed brain “total volume” which only excludes the myelencephalon and the cerebellum for reliability. In fact, the integrity of these subcortical structures heavily depends on the quality and technics used for brain extraction methods.

**Figure 5:**
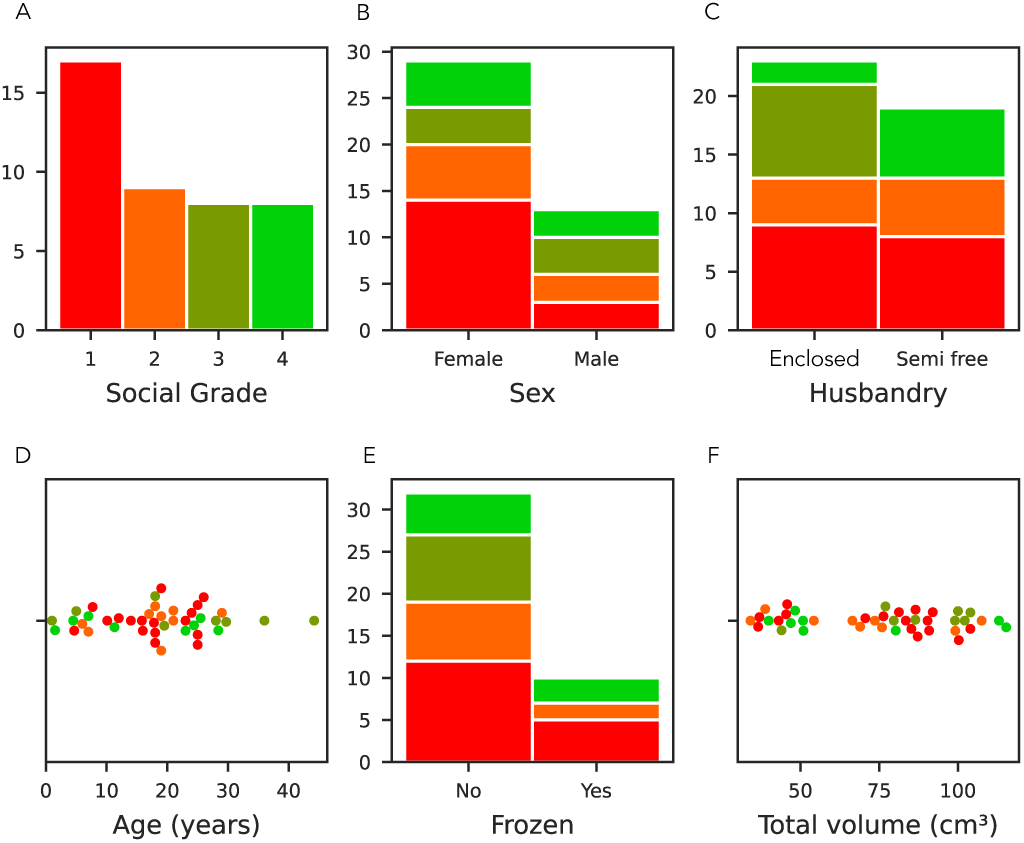
Dataset characteristics relative to the social grade. In red: social tolerance grade 1, orange: grade 2, olive: grade 3, and green: grade 4. **(A)** Social tolerance grade distribution, where grade 1 is overrepresented due to the prevalence of Macaca mulatta in laboratories. **(B)** Sex distribution: There was a significant imbalance in the sample, with females outnumbering males (2:1 ratio). **(C)** Husbandry distribution of the individuals (enclosed and semi-free ranging conditions) **(D)** Age distribution: The cohort had a relatively even age distribution with a notable peak in the 20s. **(E)** The frozen status distribution. **(F)** Total brain volume distribution, excluding the myelencephalon and cerebellum due to variation in their preservation.

### Modelling approach

To investigate the subcortical correlates of social tolerance in macaques, we used a multivariate Bayesian linear model with normal likelihood, the observed data being the amygdala and hippocampus volume. The predictors in our model were the intercept, social grade, age, sex, husbandry, whether the brain had been previously frozen, total volume, the interaction between social grade and age, and the covariance between the observations. We used wide priors, whose locations and scales were derived from the data. We assessed the quality of the model by comparing the predicted data to the observed data, and by checking the R^2^ of the model. New data was predicted to study the interaction and the age-social grade trajectory, and the difference in volume between social grades. The predictions were made using the model on all social grades, on females aged from 1 to 40, with a total volume of 85 cm^3^.

## Acknowledgement

The authors are grateful to the University of Strasbourg, the CNRS and Silabe (https://silabe.com/) for supporting this research and providing expert animal care. This work was further funded by ANR-21-CE37-0016 to S.B and J. S. This work is co funded by the French State Region contract CPER I2MT (2014-2021), R-IRM (2021-2027) and by the European Union through the European Regional Development Fund “FEDER Grand Est”. This work was performed on IRIS platform of ICube lab, member of France Life Imaging network (grant ANR 11 INBS 0006). We extend our gratitude to the PRIME-DE open science initiative, particularly the Utrecht database, as well as the Japan Monkey Center for providing access to their open science resources. Warm thanks are extended to Brice Lefaux and his staff at Mulhouse Zoo for the collection of brain samples being made possible. Additionally, we sincerely thank Aurore de Cauwer (from ICube) for her invaluable assistance at the early stages in the MRI data acquisition. The Wellcome Centre for Integrative Neuroimaging is supported by core funding from the Wellcome Trust (203139/Z/16/Z).

## Author’s contribution

S.B. and S.S. conceptualized the research; C.P., S.B., J.S. and S.S. developed the methodology, S.S. performed the data curation, J.L. performed the formal analysis, S.S., S.B. C.P., J.S. and M.L. performed the investigation; S.S., J.L. and S.B. wrote the original manuscript; S.B. and J.S. acquired the funding. All the authors reviewed the manuscript.

## Data availability

The data associated with this study are available at DOI : 10.17605/OSF.IO/AQMSW) or at this link https://doi.org/10.17605/OSF.IO/AQMSW.

## Supplemental information

Table S1: Species and data collection centers in the dataset.

Document S1: Detailed brain extraction procedure. Table S2, Figures S1-S2.

Table S3: Information relating to the acquisition of anatomical MRI images and the procedures for fixing and preserving *post-mortem* samples according to the different institutes.

## Supplementary

**Figure S1.**
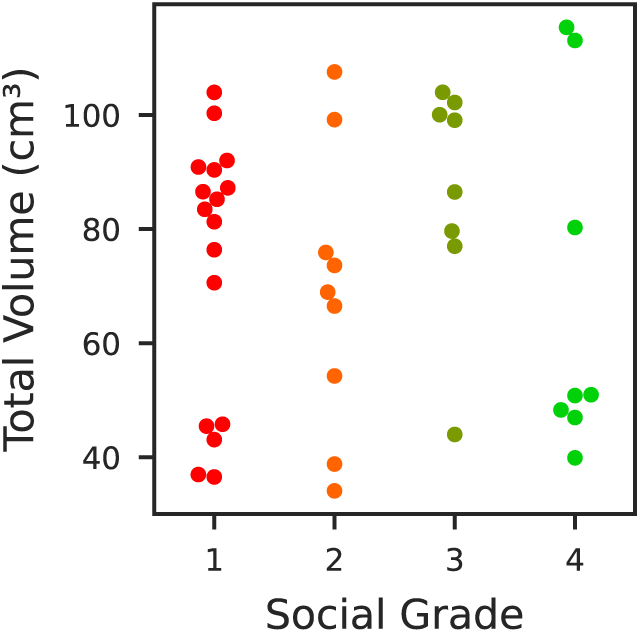
Total brain volume across macaque species categorized by social grade. Distribution of total brain volumes (in cm³) across the four social tolerance grades of the *Macaca* genus. Each dot represents an individual (n=42), and colors indicate social grade: red (grade 1, intolerant), orange (grade 2), olive (grade 3), and green (grade 4, tolerant). Total brain volume was computed from *post-mortem* MRI scans, excluding the cerebellum and myelencephalon to control for inter-individual variation in preservation quality. While total volume was included as a covariate in the statistical model, this figure provides a complementary descriptive overview of its distribution across social grades.

**Table S1:** Species and data collection centers in the dataset.

|  | CdP | Utrecht University | Inserm Oxford University | Japan Monkey Center | Total |
| --- | --- | --- | --- | --- | --- |
| *M. arctoides* [3] |  | 3 |  |  | 3 |
| *M. fascicularis* [2] | 3 | 1 | 1 | 1 | 6 |
| *M. fuscata* [1] |  | 2 |  | 2 | 4 |
| *M. mulatta* [1] | 8 | 1 | 4 |  | 13 |
| *M. nemestrina* [2] | 1 | 1 |  |  | 2 |
| *M. nigra* [4] | 1 | 1 |  |  | 2 |
| *M. silenus* [3] |  | 1 |  |  | 1 |
| *M. sinica* [3] |  |  |  | 1 | 1 |
| *M. sylvanus* [3] |  | 2 |  |  | 2 |
| *M. thibetana* [2] | 1 |  |  |  | 1 |
| *M. tonkeana* [4] | 6 |  |  |  | 6 |
| *M. radiata* [3] |  |  |  | 1 | 1 |
| Total | 20 | 12 | 5 | 5 | 42 |

Summary table of our dataset detailing the macaque species, the social grades [x] and the centers they originate from.

The dataset includes brain imaging data from 42 individuals across 12 macaque species. Columns represent the collection centers: CdP (Centre de Primatologie de l’Université de Strasbourg, France), Utrecht University (Netherlands), INSERM-Oxford University (Fr/UK), and Japan Monkey Center (Japan). Each row corresponds to a macaque species, with the number of individuals collected from each center. The total count per species and per center is indicated in the final column and row, respectively. The social grade of the species is indicated between brackets.

### Document S1: Detailed brain extraction procedure

The first part of the acquisition of brain MRI images relies on the extraction of brain specimens. All the required materials are listed in **Table S1** and have been curated through literature recommendations (Brown et al., 2009; Davenport et al., 2014; King et al., 2013; Shatil et al., 2016) as well as through the realization of the procedure itself.

**Table S2:** Necessary materials for NHP brain extraction– name and description.

| Materials | Description |
| --- | --- |
| Multifunctional rotary tool (Dremel type) associated with a sawing disc | To precisely cut out and remove the skull and have access to the brain. |
| Osteotome | To separate the skull from the atlas. |
| Mallet | Used with the osteotome. |
| Hammer-axe | To separate the skull from the atlas and to enlarge the space between the muscular planes and the skull. |
| Autopsy knife | To cut out the flesh and muscles. |
| Scalpel handle and blade | To dissect the scalp, muscles of the head, the dura, and the facial nerves. |
| Grooved probe | To dissect the dura without damaging the underlying tissues. |
| Pair of curved scissors with round tip | To cut out flesh at various steps of the dissection. |
| Dissecting forceps with and without claws | To hold tissues without damaging them to perform sections (skin, muscles, dura, nerves). |
| Graded beakers | To measure and transfer formaldehyde into the bucket for the brain fixation. |
| Funnel | To transfer the various liquids without spilling. |
| Test tube with graduation | To measure out the required amount of distilled water for the PBS. |
| Buckets (x4) | Containers for the brain fixation. |
| Hemispheric receptacles | With holes at regular intervals, to hold the brain spherical shape whilst it is immersed in formaldehyde. |
| Plastic jar with screw lid | Transparent, 7 centimeters in diameter (adapted to the size of the MRI antenna used). Container for brain during MRI acquisitions. |
| Fluorinert™ FC-770 | Fluorocarbon-based fluid, helps to displace air bubbles<sup>[1]</sup> and improves brain/background image delineation. |
| Distilled water | To dissolve the PBS tabs to make PBS solution. |
| Parafilm | To seal the MRI containers and prevent any leakage. |
| Cotton batting/aquarium foam squares | To place inside the MRI container at the bottom, the top and on one side. Used to pad the brain specimen and contain the bubble at the top of the container. |
| FFP2 mask | Safety measures for zoonosis. |
| Latex or nitril surgical gloves | For handling specimen brains. |
| Lab coats (or surgical gowns) | For individual protection whilst performing the dissection. |
| Marker | To label the various containers (mentioning if it has contained formaldehyde), label the brain specimen. |
| 7T MRI Antenna | For the MRI acquisition (whole brain acquisition). Check the internal diameter to choose the adequate container. |
| Material for the use of paraformaldehyde |  |
| Formaldehyde solution 4 % | pH 6,9 solution for histological tissue fixation. Ordered from Sigma-Aldrich (Product ID: HT501640-19L).]<br>At least 10 x volume of a macaque brain (roughly 3 liters per brain). |
| Formaldehyde spill response kits | Used for safety in case of any formaldehyde spilling. |
| Fume hood | To manipulate formaldehyde ensuring the safety of the experimenter. |
| Face shield | For the experimenter's safety in case of any spilling. |
| Mask | For the experimenter's safety in case of any spilling. |
| Gloves (chemical resistant) | For the experimenter's safety for handling the formaldehyde. |
| Baritainer | For discarding waste |

<sup>[1]</sup> (Iglesias et al., 2018)
Summary table of the tools required for the brain extraction we performed, including their
names and descriptions.

As it was the first time at the CdP that brain specimens were extracted, the development of a refined brain extraction technique played a central role in optimizing the quality of the acquired brain images. This technique, based on the literature, was meticulously applied to 17 *post-mortem* anatomical and DTI (if the specimen was not frozen) MRI scans. We optimally prepared through familiarizing with the technique and handling instruments such as the oscillating saw with a first brain extraction of a poor conditioned brain specimen (bad freezing condition). Through this first trial, we were able to adjust the following procedure as well as the chosen cutting landmarks. The brain extraction sequence consists of the following steps:

**Head Dislocation:** The animal is in supine position, with its head pulled dorsally to extend the neck (Brown et al., 2009). Then, the muscle mass of the neck has to be removed to expose the atlantooccipital junction. The junction is then incised, employing a back-and-forth motion to enable the passage of a knife or hammer-axis between the cartilage (Brown et al., 2009).

**Adjustment for Frozen Specimens:** For frozen specimens, a distinct approach through the atlantooccipital junction must be adopted. The section is performed with an osteotome and hammer **(Figure S1, A)**. Subsequently, confirmation of the absence of cerebellar protrusion through the foramen magnum is required and essential for ensuring the integrity of the specimens. If not, it would indicate cerebellar herniation and the presence of associated lesions in this brain structure (Davenport et al., 2014).

The head circumference was measured using a measuring tape during data collection. The average circumference of a macaque brain in our dataset is 30.5 ± 8.7 cm. We determined the amount of formaldehyde required based on the average volume of a macaque brain. This average volume is of 89.2 ± 1.9 (SEM) for male individuals and of 70.8 ± 0.72 cm^3^ for female individuals (Franklin et al., 2000; Scott et al., 2016). The amount of 4 % formaldehyde buffered solution (pH = 6.9, Sigma-Aldrich) required to fix the brain should follow the volume ratio of tissue to be fixed to formaldehyde of 1:10 (Thavarajah et al., 2012), that is, a minimum of one liter of formaldehyde.

**Skull Cap Removal:** The removal of the skull cap is the most meticulous part of the procedure aimed at providing access to the brain. This process starts with a longitudinal incision of the scalp from the anterior fontanel cranial suture to the foramen magnum. A second incision is made perpendicular to the first, along the coronal suture. The exposed skull surface underwent thorough cleaning with 70% ethanol, followed by drying with gauze pads. The bony skull cap was then delicately excised using a rotary tool **(Figure S1. B)**.

**Brain Extraction:** The extraction of the brain from the skull was conducted with care especially for fresh specimens as the tissues are very soft and breakable. Removal of the dura mater, cerebellum tent, and false brain is performed using a bony cap (Brown et al., 2009). Afterwards, the head is positioned vertically and some gentle taps on the table to facilitate the gradual detachment of the brain from the skull. Employing a fluted probe, the olfactory peduncles, internal carotid arteries, and cranial nerves are delicately severed (King et al., 2013). The pineal gland is commonly found as a single white firm conical mass suspended at the midline of the skull cap and attached to the meninges (King et al., 2013). In addition, in many animals, the choroid plexus is visible as two reddish masses in a position similar to or slightly in front of the pineal gland (King et al., 2013). Once the brain is fully extracted, it is placed in one of the hemispheric cups. Frozen specimens are thawed in water at room temperature before placing them in the cup (frozen tissue combined with formaldehyde fixation creates an aqueous insulating layer, altering fixation of the internal brain structures) **(Figure S1. D)**.

### Fixation and Brain Preparation for MRI

The fixation and preparation of brain specimens for MRI acquisitions is a crucial phase to ensure high quality images and low prevalence of artefacts. The chosen fixative was a 10 % formaldehyde buffered solution. Ensuring complete immersion of the brain in the fixative is crucial. To achieve this, a carefully calculated volume of the solution is added to a labelled container (usually around 3L). The brain is then placed within the container, submerged to guarantee full coverage. Sealing the container prevents contamination. The container remains for 7 days under the fume hood, and we check and gently stir every day to ensure the good repartition of the formaldehyde on the tissue. Once the brain is fixed, it is immersed the brain in a 0.5 L jar filled with PBS for 48 hours before MRI.

**Figure S1:**
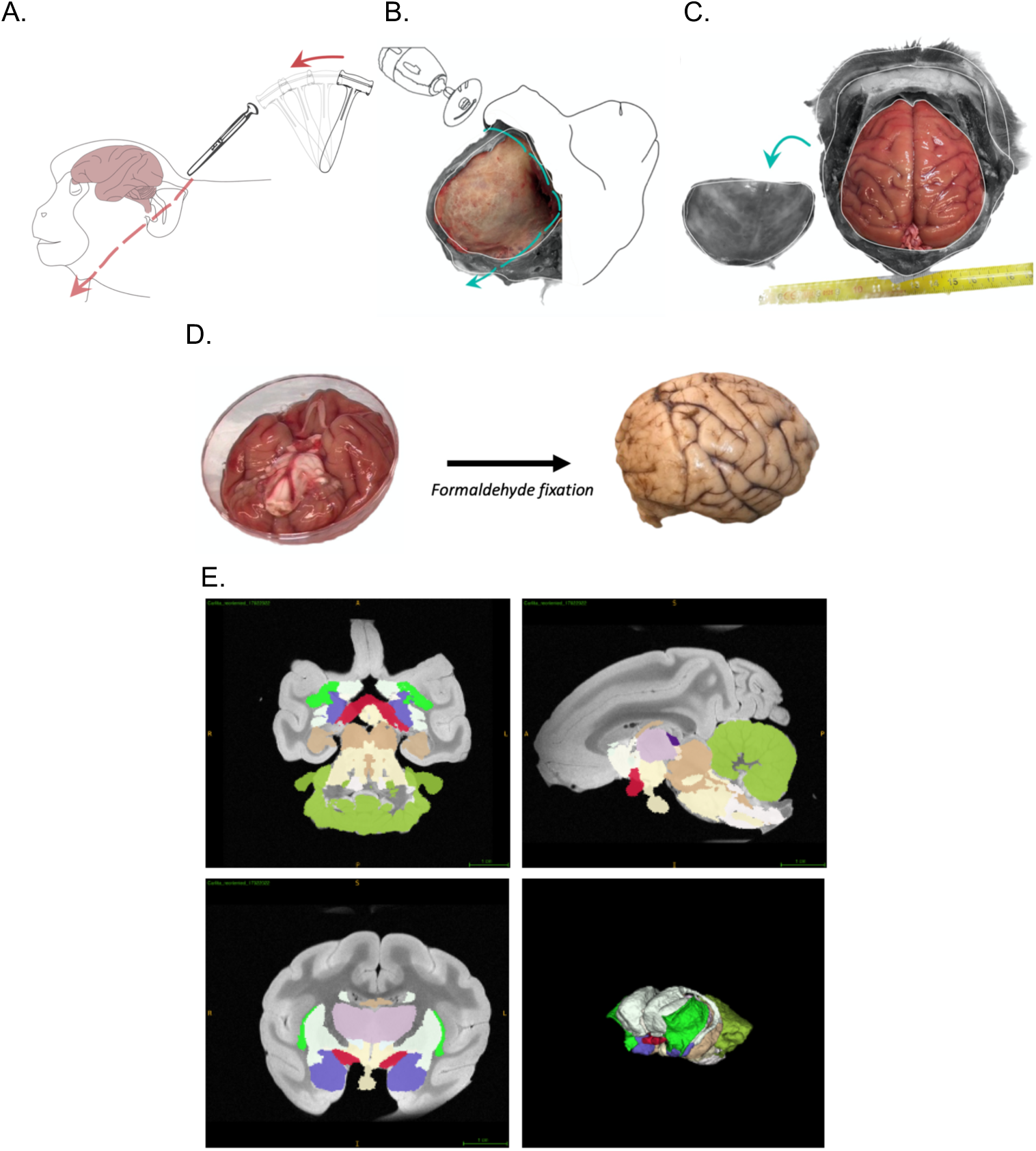
Sequence of dissection steps and MRI acquisition. (A) Craniotomy step: Position of the cadaver and cut site. (B) Steps of scalp and skull removal using a Dremel tool^®^ associated with a flex shaft rotary tool. (C) View of the skull after skull and dura removal. (D) Extraction and formaldehyde fixation of the brain. Right lateral view of the brain after a 7 day-formaldehyde-fixation. (E) SARM Regions (SARM2) and 3D MRI acquisition with atlas (bottom right). Amygdala (purple) (Hartig et al., 2021).

1. *MRI acquisitions* The diameter of the plastic container was chosen based on the diameter of the MRI antenna used (8,6 cm in diameter).

- The sample is placed in a spherical container. The orientation in which the brain is placed (for future reference during the MRI acquisitions) was registered. Aquarium foam squares are placed around the brain to minimize the residual movements from the MRI’s vibrations and to contain the remaining air bubble at the top of the container (**Figure S2. B**).
- The container is filled to the brim with Fluorinert^TM^ FC-770, a liquid that optimizes the contrast of the MRI signal and allows the wobble adjustment.
- The container is placed in a vacuum chamber (negative pressure of −0,1 Pa.), to limit the presence of air bubbles on the images, for up to three hours to remove air bubbles (Shatil et al., 2016) (Figure 1**, A**). If needed, some Fluorinert^TM^ FC-770 can be added up to the brim.
- The container is sealed with parafilm (**Figure S2. B**).
2. *Recommended 7T-MRI scanning setup* Place the container with an elevating foam square to contain the remaining bubble at the top of the jar and limit the superimposition of the bubble on the brain (**Figure S2. C and D**).

**Figure S2:**
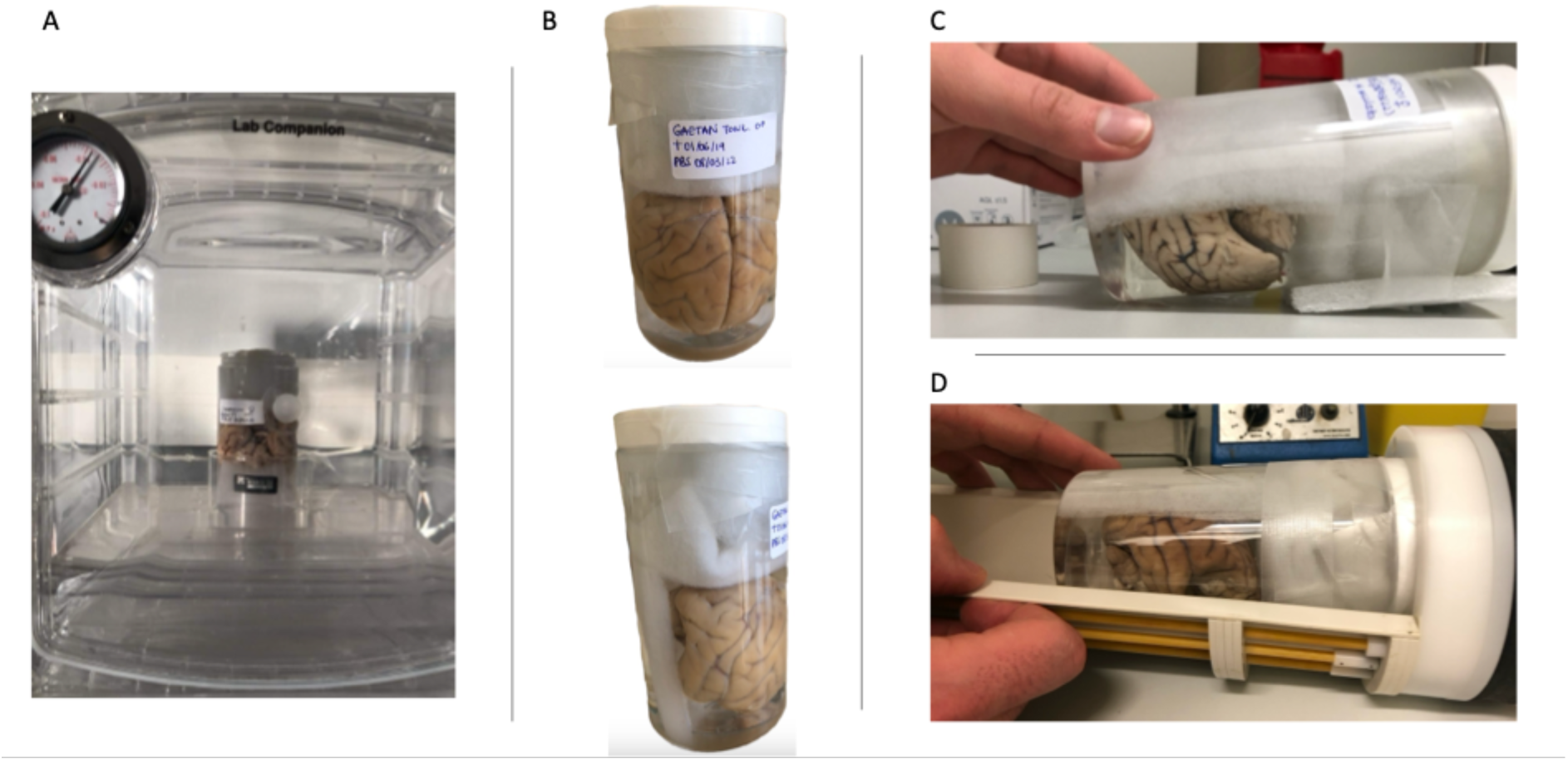
Set of photographs of the preparation of the fixed brain for MRI acquisitions. (A) Air bubble removal stages in a vacuum chamber. The brain is immersed in Fluorinert^TM^ FC-770 and held in position by the aquarium foam squares. The container is placed in a receptacle to catch any Fluorinert^TM^ FC-770 that may spill out of the container during the procedure; (B) Placement of the aquarium foam squares inside the container of brain immersed in Fluorinert^TM^ FC-770 and sealed with parafilm; (C) Placement of the container with a lift foam square to contain the residual air bubble in the top third of the container; (D) Placement of the container in the MRI antenna.

Once in the MRI, it is recommended to perform a sequence of localizer scans with the purpose of: (1) identifying significant distortions caused by air bubbles in the brain or MRI-compatible container, (2) accurately positioning the brain, and (3) establishing the slice positions necessary for subsequent data acquisition (Shatil et al., 2016).

**Table S3:** Information relating to the acquisition of anatomical MRI images and the procedures for fixing and preserving post-mortem samples according to the different centers.

| MRI Center | Primateology Center<br>(with ICube<br>laboratory) | Utrecht<br>University | Japan Monkey<br>Center | Oxford<br>University |
| --- | --- | --- | --- | --- |
| Number of<br>individuals | 19 | 14 | 4 | 4 |
| MRI | Bruker Biospec 7T<br>70/30 UR | Agilent<br>Variant 9.4T | Bruker Biospec<br>9.4T 90/20 UR | Clinical<br>Horizontal 3T |
| Antenna | Volume coil | Surface coil | Volume coil | Volume coil |
| MRI weighting | T2<br>Sequence:<br>Turbo_RARE_3D | T1 | T2 | T2 |
| Spatial<br>resolution | 0.20 x 0.20 x 0.20<br>mm | 0.25 x 0.25 x<br>0.25 mm | 0.25 x 0.25 x<br>0.25 mm | 0.6 x 0.6 x 0.6<br>mm |
| Brain fixation<br>characteristics | Immersion:<br>Formaldehyde 4% | Perfusion:<br>Formaldehyd<br>e 4% | Immersion:<br>formaline 10% | Perfusion:<br>Formaldehyde<br>4% |
| Fluid used for<br>scanning | Fluorinert® FC-770 | Sucrose<br>15%, 4°C | 3M™ Fluorinert | Fomblin or<br>Fluorinert® FC-770 |

Summary table of the MRI acquisitions parameters and *post-mortem* sample preservation protocols across centers.

## References

Adams MJ, Majolo B, Ostner J, Schülke O, De Marco A, Thierry B, Engelhardt A, Widdig A, Gerald MS, Weiss A. 2015. Personality structure and social style in macaques. J Pers Soc Psychol 109:338–353. doi:10.1037/pspp0000041

Adolphs R. 2009. The Social Brain: Neural Basis of Social Knowledge. Annu Rev Psychol 60:693–716. doi:10.1146/annurev.psych.60.110707.163514

Amaral DG, Bauman MD, Capitanio JP, Lavenex P, Mason WA, Mauldin-Jourdain ML, Mendoza SP. 2003. The amygdala: is it an essential component of the neural network for social cognition? Neuropsychologia 41:517–522. doi:10.1016/S0028-3932(02)00310-X

Amiez C, Sallet J, Giacometti C, Verstraete C, Gandaux C, Morel-Latour V, Meguerditchian A, Hadj-Bouziane F, Ben Hamed S, Hopkins WD, Procyk E, Wilson CRE, Petrides M. 2023. A revised perspective on the evolution of the lateral frontal cortex in primates. Sci Adv 9:eadf9445. doi:10.1126/sciadv.adf9445

Amiez C, Sallet J, Hopkins WD, Meguerditchian A, Hadj-Bouziane F, Ben Hamed S, Wilson CRE, Procyk E, Petrides M. 2019. Sulcal organization in the medial frontal cortex provides insights into primate brain evolution. Nat Commun 10:3437. doi:10.1038/s41467-019-11347-x

Avants BB, Tustison NJ, Song G, Cook PA, Klein A, Gee JC. 2011. A Reproducible Evaluation of ANTs Similarity Metric Performance in Brain Image Registration. Neuroimage 54:2033–2044. doi:10.1016/j.neuroimage.2010.09.025

Balasubramaniam KN, Beisner BA, Berman CM, De Marco A, Duboscq J, Koirala S, Majolo B, MacIntosh AJ, McFarland R, Molesti S, Ogawa H, Petit O, Schino G, Sosa S, Sueur C, Thierry B, de Waal FBM, McCowan B. 2017. The influence of phylogeny, social style, and sociodemographic factors on macaque social network structure. Am J Primatol 80:e22727. doi:10.1002/ajp.22727

Balasubramaniam KN, Dittmar K, Berman CM, Butovskaya M, Cooper MA, Majolo B, Ogawa H, Schino G, Thierry B, De Waal FBM. 2012. Hierarchical Steepness, Counter-Aggression, and Macaque Social Style Scale. American Journal of Primatology 74:915–925. doi:10.1002/ajp.22044

Beltrán Francés V, Castellano-Navarro A, Illa Maulany R, Ngakan PO, MacIntosh AJJ, Llorente M, Amici F. 2020. Play behavior in immature moor macaques (Macaca maura) and Japanese macaques (Macaca fuscata). American Journal of Primatology 82:e23192. doi:10.1002/ajp.23192

Berman C, Thierry B. 2010. Variation in kin bias: species differences and time constraints in macaques. doi:10.1163/000579510X539691

Bickart KC, Dickerson BC, Feldman Barrett L. 2014. The amygdala as a hub in brain networks that support social life. Neuropsychologia 63:235–248. doi:10.1016/j.neuropsychologia.2014.08.013

Bickart KC, Wright CI, Dautoff RJ, Dickerson BC, Barrett LF. 2011. Amygdala volume and social network size in humans. Nat Neurosci 14:163–164. doi:10.1038/nn.2724

Bouret S, Paradis E, Prat S, Castro L, Perez P, Gilissen E, Garcia C. 2024. Linking the evolution of two prefrontal brain regions to social and foraging challenges in primates. eLife 12:RP87780. doi:10.7554/eLife.87780

Caetano I, Amorim L, Soares JM, Ferreira S, Coelho A, Reis J, Santos NC, Moreira PS, Marques P, Magalhães R, Esteves M, Picó-Pérez M, Sousa N. 2021. Amygdala size varies with stress perception. Neurobiol Stress 14:100334. doi:10.1016/j.ynstr.2021.100334

Carlo CN, Stefanacci L, Semendeferi K, Stevens CF. 2010. Comparative analyses of the neuron numbers and volumes of the amygdaloid complex in old and new world primates. Journal of Comparative Neurology 518:1176–1198. 10.1002/cne.22264

Cooper EB, Brent LJ, Snyder-Mackler N, Singh M, Sengupta A, Khatiwada S, Malaivijitnond S, Qi Hai Z, Higham JP. 2022. The rhesus macaque as a success story of the Anthropocene. eLife 11:e78169. doi:10.7554/eLife.78169

Coplan JD, Fathy HM, Jackowski AP, Tang CY, Perera TD, Mathew SJ, Martinez J, Abdallah CG, Dwork AJ, Pantol G, Carpenter D, Gorman JM, Nemeroff CB, Owens MJ, Kaffman A, Kaufman J. 2014. Early life stress and macaque amygdala hypertrophy: preliminary evidence for a role for the serotonin transporter gene. Front Behav Neurosci 8. 10.3389/fnbeh.2014.00342

Cords M. 2012. The Behavior, Ecology, and Social Evolution of Cercopithecine Monkeys In: Mitani JC, Call J, Kappeler PM, Palombit RA, Silk JB, editors. The Evolution of Primate Societies. University of Chicago Press. pp. 91–112.

Davidson RJ, McEwen BS. 2012. Social influences on neuroplasticity: Stress and interventions to promote well-being. Nat Neurosci 15:689–695. doi:10.1038/nn.3093

de Vries H, Stevens JMG, Vervaecke H. 2006. Measuring and testing the steepness of dominance hierarchies. Animal Behaviour 71:585–592. doi:10.1016/j.anbehav.2005.05.015

De Waal FB, Johanowicz DL. 1993. Modification of Reconciliation Behavior Through Social Experience: An Experiment with Two Macaque Species. Child Development 64:897–908. doi:10.2307/1131225

De Waal FB, Luttrell LM. 1985. The formal hierarchy of rhesus macaques: An investigation of the bared-teeth display. American Journal of Primatology 9:73–85. doi:10.1002/ajp.1350090202

De Waal FBM, Luttrell LM. 1989. Toward a comparative socioecology of the genus Macaca: Different dominance styles in rhesus and stumptail monkeys. Am J Primatol 19:83–109. doi:10.1002/ajp.1350190203

Demaria, Thierry. 2001. A comparative study of reconciliation in Rhesus and Tonkean macaques. Behaviour 138:397–410. doi:10.1163/15685390152032514

DeTroy SE, Haun DBM, van Leeuwen EJC. 2021. What isn’t social tolerance? The past, present, and possible future of an overused term in the field of primatology. Evolutionary anthropology. doi:10.1002/evan.21923

Dobson SD. 2012. Coevolution of Facial Expression and Social Tolerance in Macaques. American Journal of Primatology 74:229–235. doi:10.1002/ajp.21991

Domínguez-Borràs J, Vuilleumier P. 2022. Amygdala function in emotion, cognition, and behavior. Handb Clin Neurol 187:359–380. doi:10.1016/B978-0-12-823493-8.00015-8

Duboscq J, Agil M, Engelhardt A, Thierry B. 2014. The function of postconflict interactions: new prospects from the study of a tolerant species of primate. Animal Behaviour 87:107–120. doi:10.1016/j.anbehav.2013.10.018

Duboscq J, Micheletta J, Agil M, Hodges K, Thierry B, Engelhardt A. 2013. Social Tolerance in Wild Female Crested Macaques (Macaca nigra) in Tangkoko-Batuangus Nature Reserve, Sulawesi, Indonesia. Am J Primatol 75:361–375. doi:10.1002/ajp.22114

Dubuc C, Hughes KD, Cascio J, Santos LR. 2012. Social tolerance in a despotic primate: co-feeding between consortship partners in rhesus macaques. Am J Phys Anthropol 148:73–80. doi:10.1002/ajpa.22043

Dunbar RIM. 2009. The social brain hypothesis and its implications for social evolution. Ann Hum Biol 36:562–572. doi:10.1080/03014460902960289

Dunbar RIM, Shultz S. 2025. Self-control has a social role in primates, but not in other mammals or birds. Sci Rep 15:17566. doi:10.1038/s41598-025-99523-6

Fairbanks LA. 1990. Reciprocal benefits of allomothering for female vervet monkeys. Animal Behaviour 40:553–562. doi:10.1016/S0003-3472(05)80536-6

Flack JC, Girvan M, de Waal FBM, Krakauer DC. 2006. Policing stabilizes construction of social niches in primates. Nature 439:426–429. doi:10.1038/nature04326

Freeberg TM, Dunbar RIM, Ord TJ. 2012. Social complexity as a proximate and ultimate factor in communicative complexity. Philos Trans R Soc Lond B Biol Sci 367:1785–1801. doi:10.1098/rstb.2011.0213

Ghosh A, Thakur M, Singh SK, Dutta R, Sharma LK, Chandra K, Banerjee D. 2022. The Sela macaque (*Macaca selai*) is a distinct phylogenetic species that evolved from the Arunachal macaque following allopatric speciation. Molecular Phylogenetics and Evolution 174:107513. doi:10.1016/j.ympev.2022.107513

Gumert MD, Ho M-HR. 2008. The trade balance of grooming and its coordination of reciprocation and tolerance in Indonesian long-tailed macaques (Macaca fascicularis). Primates 49:176–185. doi:10.1007/s10329-008-0089-y

Haley GE, McGuire A, Berteau-Pavy D, Weiss A, Patel R, Messaoudi I, Urbanski HF, Raber J. 2012. Measures of anxiety, amygdala volumes, and hippocampal scopolamine phMRI response in elderly female rhesus macaques. Neuropharmacology 62:385–390. doi:10.1016/j.neuropharm.2011.08.014

Han M, Jiang G, Luo H, Shao Y. 2021. Neurobiological Bases of Social Networks. Front Psychol 12. doi:10.3389/fpsyg.2021.626337

Hartig R, Glen D, Jung B, Logothetis NK, Paxinos G, Garza-Villarreal EA, Messinger A, Evrard HC. 2021. The Subcortical Atlas of the Rhesus Macaque (SARM) for neuroimaging. NeuroImage 235:117996. doi:10.1016/j.neuroimage.2021.117996

Heldstab SA, Isler K, Graber SM, Schuppli C, van Schaik CP. 2022. The economics of brain size evolution in vertebrates. Curr Biol 32:R697–R708. doi:10.1016/j.cub.2022.04.096

Heuer K, Traut N, Aristide L, Alavi SF, Herbin M, Mars RB, Mylapalli R, Najafipashaki S, Sakai T, Santin M, Borrell V, Toro R. 2025. Principles of neocortical organisation and behaviour in primates. doi:10.1101/2025.07.17.665410

Hill DA, Okayasu N. 1995. Absence of “Youngest Ascendancy” in the Dominance Relations of Sisters in Wild Japanese Macaques (Macaca Fuscata Yakui). doi:10.1163/156853995X00612

Hölzel BK, Carmody J, Evans KC, Hoge EA, Dusek JA, Morgan L, Pitman RK, Lazar SW. 2010. Stress reduction correlates with structural changes in the amygdala. Social Cognitive and Affective Neuroscience 5:11–17. doi:10.1093/scan/nsp034

Howard AFD, Huszar IN, Smart A, Cottaar M, Daubney G, Hanayik T, Khrapitchev AA, Mars RB, Mollink J, Scott C, Sibson NR, Sallet J, Jbabdi S, Miller KL. 2023. An open resource combining multi-contrast MRI and microscopy in the macaque brain. Nat Commun 14:4320. doi:10.1038/s41467-023-39916-1

Howell BR, Grand AP, McCormack KM, Shi Y, LaPrarie J, Maestripieri D, Styner MA, Sanchez MM. 2014. Early Adverse Experience Increases Emotional Reactivity in Juvenile Rhesus Macaques: Relation to Amygdala Volume. Dev Psychobiol 56:1735–1746. doi:10.1002/dev.21237

Isa T, Yamane I, Hamai M, Inagaki H. 2009. Japanese macaques as laboratory animals. Exp Anim 58:451–457. doi:10.1538/expanim.58.451

Joly M, Micheletta J, De Marco A, Langermans JA, Sterck EHM, Waller BM. 2017. Comparing physical and social cognitive skills in macaque species with different degrees of social tolerance. Proc Royal Soc B 284:20162738. doi:10.1098/rspb.2016.2738

Jones DN, Erwin JM, Sherwood CC, Hof PR, Raghanti MA. 2021. A comparison of cell density and serotonergic innervation of the amygdala among four macaque species. J Comp Neurol 529:1659–1668. doi:10.1002/cne.25048

Jung B, Taylor PA, Seidlitz J, Sponheim C, Perkins P, Ungerleider LG, Glen D, Messinger A. 2021. A comprehensive macaque fMRI pipeline and hierarchical atlas. NeuroImage 235:117997. doi:10.1016/j.neuroimage.2021.117997

Kanai R, Bahrami B, Roylance R, Rees G. 2012. Online social network size is reflected in human brain structure. Proc Royal Soc B 279:1327–1334. doi:10.1098/rspb.2011.1959

Kim EJ, Pellman B, Kim JJ. 2015. Stress effects on the hippocampus: a critical review. Learn Mem 22:411–416. doi:10.1101/lm.037291.114

Kruschke JK. 2015. Doing Bayesian Data Analysis - A Tutorial with R, JAGS, and Stan, 2nd ed. Boston: Academic Press.

Kutsukake N. 2000. Matrilineal rank Inheritance varies with absolute rank in Japanese macaques. Primates 41:321–335. doi:10.1007/BF02557601

Liebal K, Waller BM, Burrows AM, Slocombe KE. 2014. Primate communication: a multimodal approach. Cambridge: Cambridge University Press.

Loyant L, Waller BM, Micheletta J, Meunier H, Ballesta S, Joly M. 2023. Tolerant macaque species are less impulsive and reactive. Anim Cogn 26:1453–1466. doi:25

Lyons DM, Parker KJ, Zeitzer JM, Buckmaster CL, Schatzberg AF. 2007. Preliminary evidence that hippocampal volumes in monkeys predict stress levels of adrenocorticotropic hormone. Biol Psychiatry 62:1171–1174. doi:10.1016/j.biopsych.2007.03.012

Lyons DM, Yang C, Sawyer-Glover AM, Moseley ME, Schatzberg AF. 2001. Early life stress and inherited variation in monkey hippocampal volumes. Arch Gen Psychiatry 58:1145–1151. doi:10.1001/archpsyc.58.12.1145

Maestripieri D. 1994. Social structure, infant handling, and mothering styles in group-living old world monkeys. Int J Primatol 15:531–553. doi:10.1007/BF02735970

Maguire EA, Gadian DG, Johnsrude IS, Good CD, Ashburner J, Frackowiak RS, Frith CD. 2000. Navigation-related structural change in the hippocampi of taxi drivers. Proc Natl Acad Sci U S A 97:4398–4403. doi:10.1073/pnas.070039597

ManyPrimates, Aguenounon G, Allritz M, Ballesta S. 2022. The evolution of primate short-term memory. Animal Behavior and Cognition 9:428–516. doi:10.26451/abc.09.04.06.2022

Maranesi M, Livi A, Fogassi L, Rizzolatti G, Bonini L. 2014. Mirror neuron activation prior to action observation in a predictable context. J Neurosci 34:14827–14832. doi:10.1523/JNEUROSCI.2705-14.2014

Massen JJM, Berg LM van den, Spruijt BM, Sterck EHM. 2010. Generous Leaders and Selfish Underdogs: Pro-Sociality in Despotic Macaques. PLOS ONE 5:e9734. doi:10.1371/journal.pone.0009734

McCowan B, Vandeleest J, Balasubramaniam K, Hsieh F, Nathman A, Beisner B. 2022. Measuring dominance certainty and assessing its impact on individual and societal health in a nonhuman primate model: a network approach. Philosophical Transactions of the Royal Society B: Biological Sciences 377:20200438. doi:10.1098/rstb.2020.0438

Meguerditchian A, Marie D, Margiotoudi K, Roth M, Nazarian B, Anton J-L, Claidière N. 2021. Baboons (Papio anubis) living in larger social groups have bigger brains. Evolution and Human Behavior 42:30–34. doi:10.1016/j.evolhumbehav.2020.06.010

Meyer JS, Hamel AF. 2014. Models of Stress in Nonhuman Primates and Their Relevance for Human Psychopathology and Endocrine Dysfunction. ILAR Journal 55:347–360. doi:10.1093/ilar/ilu023

Micheletta J, Waller BM, Panggur MR, Neumann C, Duboscq J, Agil M, Engelhardt A. 2012. Social bonds affect anti-predator behaviour in a tolerant species of macaque, Macaca nigra. Proc Biol Sci 279:4042–4050. doi:10.1098/rspb.2012.1470

Milham MP, Ai L, Koo B, Xu T, Amiez C, Balezeau F, Baxter MG, Blezer ELA, Brochier T, Chen A, Croxson PL, Damatac CG, Dehaene S, Everling S, Fair DA, Fleysher L, Freiwald W, Froudist-Walsh S, Griffiths TD, Guedj C, Hadj-Bouziane F, Ben Hamed S, Harel N, Hiba B, Jarraya B, Jung B, Kastner S, Klink PC, Kwok SC, Laland KN, Leopold DA, Lindenfors P, Mars RB, Menon RS, Messinger A, Meunier M, Mok K, Morrison JH, Nacef J, Nagy J, Rios MO, Petkov CI, Pinsk M, Poirier C, Procyk E, Rajimehr R, Reader SM, Roelfsema PR, Rudko DA, Rushworth MFS, Russ BE, Sallet J, Schmid MC, Schwiedrzik CM, Seidlitz J, Sein J, Shmuel A, Sullivan EL, Ungerleider L, Thiele A, Todorov OS, Tsao D, Wang Z, Wilson CRE, Yacoub E, Ye FQ, Zarco W, Zhou Y, Margulies DS, Schroeder CE. 2018. An Open Resource for Non-human Primate Imaging. Neuron 100:61–74.e2. doi:10.1016/j.neuron.2018.08.039

Mulholland MM, Hecht E, Wesley MJ, Hopkins WD. 2024. Long term impacts of early social environment on chimpanzee white matter. Sci Rep 14:29879. doi:10.1038/s41598-024-81238-9

Navarrete AF, Blezer ELA, Pagnotta M, de Viet ESM, Todorov O, Lindenfors P, Laland K, Reader SM. 2018. Primate Brain Anatomy: New Volumetric MRI Measurements for Neuroanatomical Studies. Brain Behav Evol 91:1–9. doi:10.1159/000488136

Noonan MP, Sallet J, Mars RB, Neubert FX, O’Reilly JX, Andersson JL, Mitchell AS, Bell AH, Miller KL, Rushworth MFS. 2014a. A Neural Circuit Covarying with Social Hierarchy in Macaques. PLoS Biology 12:e1001940. doi:10.1371/journal.pbio.1001940

Noonan MP, Sallet J, Mars RB, Neubert FX, O’Reilly JX, Andersson JL, Mitchell AS, Bell AH, Miller KL, Rushworth MFS. 2014b. A Neural Circuit Covarying with Social Hierarchy in Macaques. PLOS Biol 12:e1001940. doi:10.1371/journal.pbio.1001940

Ochsner KN, Gross JJ. 2008. Cognitive Emotion Regulation: Insights from Social Cognitive and Affective Neuroscience. Curr Dir Psychol Sci 17:153–158. doi:10.1111/j.1467-8721.2008.00566.x

Parkinson C, Kleinbaum AM, Wheatley T. 2017. Spontaneous neural encoding of social network position. Nat Hum Behav 1:1–7. doi:10.1038/s41562-017-0072

Passingham RE, Wise SP, Passingham RE, Wise SP. 2012. The Neurobiology of the Prefrontal Cortex: Anatomy, Evolution, and the Origin of Insight, Oxford Psychology Series. Oxford, New York: Oxford University Press.

Patzelt A, Kopp GH, Ndao I, Kalbitzer U, Zinner D, Fischer J. 2014. Male tolerance and male– male bonds in a multilevel primate society. Proceedings of the National Academy of Sciences 111:14740–14745. doi:10.1073/pnas.1405811111

Perelman P, Johnson WE, Roos C, Seuánez HN, Horvath JE, Moreira MAM, Kessing B, Pontius J, Roelke M, Rumpler Y, Schneider MPC, Silva A, O’Brien SJ, Pecon-Slattery J. 2011. A Molecular Phylogeny of Living Primates. PLoS Genet 7:e1001342. doi:10.1371/journal.pgen.1001342

Pessoa L. 2010. Emotion and Cognition and the Amygdala: From “what is it?” to “what’s to be done?” Neuropsychologia 48:3416–3429. doi:10.1016/j.neuropsychologia.2010.06.038

Petit O, Abegg C, Thierry B. 1997. A comparative study of aggression and conciliation in three cercopithecine monkeys (Macaca fuscata, Macaca nigra, Papio papio). Behaviour 134:415–432. doi:10.1163/156853997X00610

Petit O, Bertrand F, Thierry B. 2008. Social play in crested and Japanese macaques: testing the covariation hypothesis. Dev Psychobiol 50:399–407. doi:10.1002/dev.20305

Phelps EA. 2006. Emotion and cognition: insights from studies of the human amygdala. Annu Rev Psychol 57:27–53. doi:10.1146/annurev.psych.56.091103.070234

Phelps EA, LeDoux JE. 2005. Contributions of the amygdala to emotion processing: from animal models to human behavior. Neuron 48:175–187. doi:10.1016/j.neuron.2005.09.025

Rincon AV, Waller BM, Duboscq J, Mielke A, Pérez C, Clark PR, Micheletta J. 2023. Higher social tolerance is associated with more complex facial behavior in macaques. Elife 12:RP87008. doi:10.7554/eLife.87008

Sadoughi B, Lacroix L, Berbesque C, Meunier H, Lehmann J. 2021. Effects of social tolerance on stress: hair cortisol concentrations in the tolerant Tonkean macaques (Macaca tonkeana) and the despotic long-tailed macaques (Macaca fascicularis). Stress 24:1033–1041. doi:10.1080/10253890.2021.1998443

Sakai T, Hata J, Ohta H, Shintaku Y, Kimura N, Ogawa Y, Sogabe K, Mori S, Okano HJ, Hamada Y, Shibata S, Okano H, Oishi K. 2018. The Japan Monkey Centre Primates Brain Imaging Repository for comparative neuroscience: an archive of digital records including records for endangered species. Primates 59:553–570. doi:10.1007/s10329-018-0694-3

Sakai T, Hata J, Shintaku Y, Ohta H, Sogabe K, Mori S, Miyabe-Nishiwaki T, Okano HJ, Hamada Y, Hirabayashi T, Minamimoto T, Sadato N, Okano H, Oishi K. 2023. The Japan Monkey Centre Primates Brain Imaging Repository of high-resolution postmortem magnetic resonance imaging: The second phase of the archive of digital records. NeuroImage 273:120096. doi:10.1016/j.neuroimage.2023.120096

Sallet J, Mars RB, Noonan MP, Andersson JL, O’Reilly JX, Jbabdi S, Croxson PL, Jenkinson M, Miller KL, Rushworth MFS. 2011. Social Network Size Affects Neural Circuits in Macaques. Science 334:697–700. doi:10.1126/science.1210027

Sapolsky RM, Uno H, Rebert CS, Finch CE. 1990. Hippocampal damage associated with prolonged glucocorticoid exposure in primates. J Neurosci 10:2897–2902. doi:10.1523/JNEUROSCI.10-09-02897.1990

Schumann CM, Scott JA, Lee A, Bauman MD, Amaral DG. 2019. Amygdala growth from youth to adulthood in the macaque monkey. J Comp Neurol 527:3034–3045. doi:10.1002/cne.24728

Scopa C, Palagi E. 2016. Mimic me while playing! Social tolerance and rapid facial mimicry in macaques (Macaca tonkeana and Macaca fuscata). J Comp Psychol 130:153–161. doi:10.1037/com0000028

Sébille SB, Rolland A-S, Welter M-L, Bardinet E, Santin MD. 2019. Post mortem high resolution diffusion MRI for large specimen imaging at 11.7 T with 3D segmented echo-planar imaging. Journal of Neuroscience Methods 311:222–234. doi:10.1016/j.jneumeth.2018.10.010

Silk JB. 2002. Kin Selection in Primate Groups. International Journal of Primatology 23:849–875. doi:10.1023/A:1015581016205

Silk JB. 1999. Male bonnet macaques use information about third-party rank relationships to recruit allies. Animal Behaviour 58:45–51. doi:10.1006/anbe.1999.1129

Silk JB, Alberts SC, Altmann J. 2003. Social Bonds of Female Baboons Enhance Infant Survival. Science 302:1231–1234. doi:10.1126/science.1088580

Song X, García-Saldivar P, Kindred N, Wang Y, Merchant H, Meguerditchian A, Yang Y, Stein EA, Bradberry CW, Ben Hamed S, Jedema HP, Poirier C. 2021. Strengths and challenges of longitudinal non-human primate neuroimaging. Neuroimage 236:118009. doi:10.1016/j.neuroimage.2021.118009

Sueur C, Petit O, De Marco A, Jacobs AT, Watanabe K, Thierry B. 2011. A comparative network analysis of social style in macaques. Animal Behaviour 82:845–852. doi:10.1016/j.anbehav.2011.07.020

Suomi SJ. 2008. Attachment in rhesus monkeysHandbook of Attachment: Theory, Research, and Clinical Applications, 2nd Ed. New York, NY, US: The Guilford Press. pp. 173–191.

Taren AA, Creswell JD, Gianaros PJ. 2013. Dispositional Mindfulness Co-Varies with Smaller Amygdala and Caudate Volumes in Community Adults. PLoS One 8:e64574. doi:10.1371/journal.pone.0064574

Testard C, Brent LJN, Andersson J, Chiou KL, Negron-Del Valle JE, DeCasien AR, Acevedo-Ithier A, Stock MK, Antón SC, Gonzalez O, Walker CS, Foxley S, Compo NR, Bauman S, Ruiz-Lambides AV, Martinez MI, Skene JHP, Horvath JE, Unit CBR, Higham JP, Miller KL, Snyder-Mackler N, Montague MJ, Platt ML, Sallet J. 2022. Social connections predict brain structure in a multidimensional free-ranging primate society. Sci Adv 8:eabl5794. doi:10.1126/sciadv.abl5794

Thierry B. 2021. Where do we stand with the covariation framework in primate societies? Am J Biol Anthropol 178. doi:10.1002/ajpa.24441

Thierry B. 2017. Macaque (Macaca)The International Encyclopedia of Primatology. John Wiley & Sons, Ltd. pp. 1–4. doi:10.1002/9781119179313.wbprim0040

Thierry B. 2007. Unity in diversity: Lessons from macaque societies. Evol Anthropol 16:224–238. doi:10.1002/evan.20147

Thierry B. 2000. Covariation of Conflict Management Patterns across Macaque SpeciesNatural Conflict Resolution. Berkeley, CA, US: University of California Press. pp. 106–128.

Thierry B, Aureli F, Nunn CL, Petit O, Abegg C, de Waal FBM. 2008. A comparative study of conflict resolution in macaques: insights into the nature of trait covariation. Animal Behaviour 75:847–860. doi:10.1016/j.anbehav.2007.07.006

Thierry B, Sapolsky R. 2000. CHAPTER 6 Covariation of Conflict Management Patterns across Macaque Species In: Aureli F, editor. Natural Conflict Resolution. University of California Press. pp. 106–128.

Thierry B, Singh M, Kaumanns W. 2004. Macaque Societies: A Model for the Study of Social Organization. Cambridge University Press.

Todorov OS, Weisbecker V, Gilissen E, Zilles K, de Sousa AA. 2019. Primate hippocampus size and organization are predicted by sociality but not diet. Proceedings of the Royal Society B: Biological Sciences 286:20191712. doi:10.1098/rspb.2019.1712

Tottenham N, Hare TA, Quinn BT, McCarry TW, Nurse M, Gilhooly T, Millner A, Galvan A, Davidson MC, Eigsti I-M, Thomas KM, Freed PJ, Booma ES, Gunnar MR, Altemus M, Aronson J, Casey B j. 2010. Prolonged institutional rearing is associated with atypically large amygdala volume and difficulties in emotion regulation. Developmental Science 13:46–61. doi:10.1111/j.1467-7687.2009.00852.x

Uematsu A, Matsui M, Tanaka C, Takahashi T, Noguchi K, Suzuki M, Nishijo H. 2012. Developmental Trajectories of Amygdala and Hippocampus from Infancy to Early Adulthood in Healthy Individuals. PLoS One 7:e46970. doi:10.1371/journal.pone.0046970

Vandeleest JJ, Beisner BA, Hannibal DL, Nathman AC, Capitanio JP, Hsieh F, Atwill ER, McCowan B. 2016. Decoupling social status and status certainty effects on health in macaques: a network approach. PeerJ 4:e2394. doi:10.7717/peerj.2394

Waller BM, Liebal K, Burrows AM, Slocombe Katie E. 2013. How Can a Multimodal Approach to Primate Communication Help Us Understand the Evolution of Communication? Evol Psychol 11:538–549. doi:10.1177/147470491301100305

Waller BM, Whitehouse J, Micheletta J. 2016. Macaques can predict social outcomes from facial expressions. Anim Cogn 19:1031–1036. doi:10.1007/s10071-016-0992-3

Whitehouse J, Clark PR, Robinson RL, Rees K, O’Callaghan O, Kimock CM, Witham CL, Waller BM. 2024. Facial expressivity in dominant macaques is linked to group cohesion. Proc Biol Sci 291:20240984. doi:10.1098/rspb.2024.0984

Yushkevich PA, Piven J, Hazlett HC, Smith RG, Ho S, Gee JC, Gerig G. 2006. User-guided 3D active contour segmentation of anatomical structures: Significantly improved efficiency and reliability. NeuroImage 31:1116–1128. doi:10.1016/j.neuroimage.2006.01.015

Zannella A, Stanyon R, Palagi E. 2017. Yawning and social styles: Different functions in tolerant and despotic macaques (Macaca tonkeana and Macaca fuscata). Journal of Comparative Psychology 131:179–188. doi:10.1037/com0000062

Zhang D, Guo L, Zhu D, Li K, Li L, Chen H, Zhao Q, Hu X, Liu T. 2013. Diffusion Tensor Imaging Reveals Evolution of Primate Brain Architectures. Brain Struct Funct 218:10.1007/s00429-012-0468–4. doi:10.1007/s00429-012-0468-4

Zou KH, Warfield SK, Bharatha A, Tempany CMC, Kaus MR, Haker SJ, Wells WM, Jolesz FA, Kikinis R. 2004. Statistical Validation of Image Segmentation Quality Based on a Spatial Overlap Index. Acad Radiol 11:178–189. doi:10.1016/S1076-6332(03)00671-8

## Supplementary references

Brown C, Rech RR, Torres F. 2009. A Field Manual for Collection of Specimens to Enhance Diagnosis of Animal Diseases. Boca Raton, FL, USA: Boca Publications Group, Incorporated.

Davenport AT, Grant KA, Szeliga KT, Friedman DP, Daunais JB. 2014. Standardized Method for the Harvest of Nonhuman Primate Tissue Optimized for Multiple Modes of Analyses. Cell Tissue Bank 15:99–110. doi:10.1007/s10561-013-9380-2

Franklin MS, Kraemer GW, Shelton SE, Baker E, Kalin NH, Uno H. 2000. Gender differences in brain volume and size of corpus callosum and amygdala of rhesus monkey measured from MRI images. Brain Res 852:263–267. doi:10.1016/s0006-8993(99)02093-4

Iglesias JE, Crampsie S, Strand C, Tachrount M, Thomas DL, Holton JL. 2018. Effect of Fluorinert on the Histological Properties of Formalin-Fixed Human Brain Tissue. J Neuropathol Exp Neurol 77:1085–1090. doi:10.1093/jnen/nly098

King JM, Roth-Johnson L, Dodd DC, Newsom ME. 2013. The Necropsy Book: A Guide for Veterinary Students, Residents, Clinicians, Pathologists and Biological Researchers, 7th ed. Cornell University, Ithaca, New York, 14850: College of Veterinary Medicine.

Scott JA, Grayson D, Fletcher E, Lee A, Bauman MD, Schumann CM, Buonocore MH, Amaral DG. 2016. Longitudinal analysis of the developing rhesus monkey brain using magnetic resonance imaging: birth to adulthood. Brain Struct Funct 221:2847–2871. doi:10.1007/s00429-015-1076-x

Shatil AS, Matsuda KM, Figley CR. 2016. A Method for Whole Brain Ex Vivo Magnetic Resonance Imaging with Minimal Susceptibility Artifacts. Frontiers in Neurology 7.

Thavarajah R, Mudimbaimannar VK, Elizabeth J, Rao UK, Ranganathan K. 2012. Chemical and physical basics of routine formaldehyde fixation. J Oral Maxillofac Pathol 16:400–405. doi:10.4103/0973-029X.102496

